# A robust brain network for sustained attention from adolescence to adulthood that predicts later substance use

**DOI:** 10.1101/2024.04.03.587900

**Authors:** Yihe Weng, Johann Kruschwitz, Laura M. Rueda-Delgado, Kathy Ruddy, Rory Boyle, Luisa Franzen, Emin Serin, Tochukwu Nweze, Jamie Hanson, Alannah Smyth, Tom Farnan, Tobias Banaschewski, Arun L.W. Bokde, Sylvane Desrivières, Herta Flor, Antoine Grigis, Hugh Garavan, Penny Gowland, Andreas Heinz, Rüdiger Brühl, Jean-Luc Martinot, Marie-Laure Paillère Martinot, Eric Artiges, Jane McGrath, Frauke Nees, Dimitri Papadopoulos Orfanos, Tomáš Paus, Luise Poustka, Nathalie Holz, Juliane H. Fröhner, Michael N. Smolka, Nilakshi Vaidya, Gunter Schumann, Henrik Walter, Robert Whelan, IMAGEN Consortium

**Affiliations:** School of Psychology and Global Brain Health Institute, Trinity College Dublin, Ireland; Department of Psychiatry and Psychotherapy CCM, Charité – Universitätsmedizin Berlin, corporate member of Freie Universität Berlin, Humboldt-Universität zu Berlin, and Berlin Institute of Health, Berlin, Germany; Collaborative Research Centre (SFB 940) “Volition and Cognitive Control”, Technische Universität Dresden, 01069, Dresden, Germany; School of Psychology, Queens University Belfast, Belfast, Northern Ireland, UK; Faculty of Psychology and Neuroscience, Maastricht University, Maastricht, Netherlands; Charité –Universitätsmedizin Berlin, Einstein Center for Neurosciences Berlin, 10117, Berlin, Germany; Bernstein Center for Computational Neuroscience, 10115, Berlin, Germany; Department of Psychology, University of Utah, USA; Department of Psychology, University of Pittsburgh, Pittsburgh, PA, USA; Learning Research & Development Center, University of Pittsburgh, Pittsburgh, PA, USA; Department of Child and Adolescent Psychiatry and Psychotherapy, Central Institute of Mental Health, Medical Faculty Mannheim, Heidelberg University, Square J5, 68159 Mannheim, Germany; Discipline of Psychiatry, School of Medicine and Trinity College Institute of Neuroscience, Trinity College Dublin, Dublin, Ireland; Centre for Population Neuroscience and Precision Medicine (PONS), Institute of Psychiatry, Psychology & Neuroscience, SGDP Centre, King’s College London, United Kingdom; Institute of Cognitive and Clinical Neuroscience, Central Institute of Mental Health, Medical Faculty Mannheim, Heidelberg University, Square J5, Mannheim, Germany; Department of Psychology, School of Social Sciences, University of Mannheim, 68131 Mannheim, Germany; NeuroSpin, CEA, Université Paris-Saclay, F-91191 Gif-sur-Yvette, France; Departments of Psychiatry and Psychology, University of Vermont, 05405 Burlington, Vermont, USA; Sir Peter Mansfield Imaging Centre School of Physics and Astronomy, University of Nottingham, University Park, Nottingham, United Kingdom; Physikalisch-Technische Bundesanstalt (PTB), Braunschweig and Berlin, Germany; Institut National de la Santé et de la Recherche Médicale, INSERM U 1299 “Trajectoires développementales & psychiatrie”, University Paris-Saclay, CNRS; Ecole Normale Supérieure Paris-Saclay, Centre Borelli; Gif-sur-Yvette, France; Institut National de la Santé et de la Recherche Médicale, INSERM U 1299 “Trajectoires développementales & psychiatrie”, University Paris-Saclay, CNRS; Ecole Normale Supérieure Paris-Saclay, Centre Borelli; Gif-sur-Yvette; and AP-HP. Sorbonne University, Department of Child and Adolescent Psychiatry, Pitié-Salpêtrière Hospital, Paris, France; Institut National de la Santé et de la Recherche Médicale, INSERM U 1299 “Trajectoires développementales & psychiatrie”, University Paris-Saclay, CNRS; Ecole Normale Supérieure Paris-Saclay, Centre Borelli; Gif-sur-Yvette; and Psychiatry Department, EPS Barthélémy Durand, Etampes, France; Institute of Medical Psychology and Medical Sociology, University Medical Center Schleswig Holstein, Kiel University, Kiel, Germany; Departments of Psychiatry and Neuroscience, Faculty of Medicine and Centre Hosptalier UniversitaireSainte-Justine, University of Montreal, Montreal, Quebec, Canada; Departments of Psychiatry and Psychology, University of Toronto, Toronto, Ontario, Canada; Department of Child and Adolescent Psychiatry and Psychotherapy, University Medical Centre Göttingen, von-Siebold-Str. 5, 37075, Göttingen, Germany; Department of Psychiatry and Neuroimaging Center, Technische Universität Dresden, Dresden, Germany; Centre for Population Neuroscience and Stratified Medicine (PONS), Department of Psychiatry and Neuroscience, Charité Universitätsmedizin Berlin, Germany; Centre for Population Neuroscience and Precision Medicine (PONS), Institute for Science and Technology of Brain-inspired Intelligence (ISTBI), Fudan University, Shanghai, China

## Abstract

Substance use, including cigarettes and cannabis, is associated with poorer sustained attention in late adolescence and early adulthood. Previous studies were predominantly cross-sectional or under-powered and could not indicate if impairment in sustained attention was a predictor of substance-use or a marker of the inclination to engage in such behaviour. This study explored the relationship between sustained attention and substance use across a longitudinal span from ages 14 to 23 in over 1,000 participants. Behaviours and brain connectivity associated with diminished sustained attention at age 14 predicted subsequent increases in cannabis and cigarette smoking, establishing sustained attention as a robust biomarker for vulnerability to substance use. Individual differences in network strength relevant to sustained attention were preserved across developmental stages and sustained attention networks generalized to participants in an external dataset. In summary, brain networks of sustained attention are robust, consistent, and able to predict aspects of later substance use.

**Teaser:** A robust brain network for sustained attention at age 14 predicts cigarette and cannabis use from ages 14 to 23.

## Introduction

Sustained attention is a critical cognitive process in daily life, playing a significant role in academic achievement, social communication, and mental health (Esterman and Rothlein, 2019) and can be defined as “the focus on performance on a single task over time, with the goal of explaining both the fluctuations within an individual as well as the individual differences in overall ability to maintain stable task performance” (p. 174)(Esterman and Rothlein, 2019). Sustained attention notably improves between the ages of 9 and 16 years-old (Thomson et al., 2022), concomitant with cognitive maturation and brain development during adolescence (Paus, 2005). The functional neuroanatomy of sustained attention involves cingulate, prefrontal, and parietal cortices; supplementary motor area (SMA); frontal eye field; and cerebellum (Bauer et al., 2020; Pinar et al., 2018).

Cross-sectional studies suggest that substance use during adolescence, including cigarette smoking (Treur et al., 2015), alcohol consumption (Ueno et al., 2022), and cannabis use (Wallace et al., 2019), is associated with poorer sustained attention. For instance, adolescents (14-17 years-old) using cannabis a minimum of 4 days per week for at least the last 6 months showed impaired sustained attention in the Rapid Visual Information Processing Task (RVP), and in the Immediate Memory Task versus non-users (Dougherty et al., 2013). Adolescents (12-17 years-old) in a high tetrahydrocannabinol (THC; the primary psychoactive component in cannabis) group exhibited lower accuracy on the RVP task than a low THC group (Shannon et al., 2010). Cigarette users aged 18-29 years-old showed significant cognitive impairments in sustained attention than non-smokers in the RVP task (Chamberlain et al., 2012). A systematic review of the next-day cognitive effects of heavy alcohol consumption demonstrated impairments in sustained attention during alcohol hangovers using meta-analysis (Yakir et al., 2007). These findings highlight the negative associations between substance use and sustained attention.

Given the cross-sectional nature of the behavioural and neuroimaging studies above, it remains unclear if impaired sustained attention predates the initiation of substance use and/or if it is a consequence of substance use. Only one longitudinal study (Harakeh et al., 2012) has examined the association between sustained attention and cigarette smoking, employing measurements across three-waves and involving a large sample of 1,797 adolescents. Poor sustained attention, unlike other neurocognitive functions such as working memory, attention flexibility, or perceptual sensitivity; was associated with the increased probability of adolescents subsequently initiating cigarette smoking between ages 11 and 13 and with a higher chance of being a daily smoker by age 16. Harakeh and colleagues’ findings suggest that poor sustained attention may precede the onset of cigarette smoking. However, as their study was based on a behavioural level, the neural correlates underlying these associations remain untested.

Although lower sustained attention has been associated with subsequent cigarette smoking, individuals commonly engage in the concurrent use of multiple substances (Crummy et al., 2020), perhaps due to shared pathological substrates for substance use. A meta-analysis identified common neural alterations in primary dorsal striatal, and frontal circuits, engaged in reward/salience processing, habit formation, and executive control across various substances (nicotine, cannabis, alcohol, and cocaine) (Thiele and Bellgrove, 2018). Those involved in substance use often co-use both cannabis and cigarettes (Agrawal et al., 2012; Hindocha et al., 2016; Weinberger et al., 2018). Agrawal et al. (2012) reported that 90% of cannabis users smoke cigarettes during their lifetime, and the widespread co-use of the two may be attributed to genetic sharing (Agrawal et al., 2010; Stringer et al., 2016) and similar neural mechanisms (Klugah-Brown et al., 2020).

Functional brain networks can predict various behavioural traits, such as substance use (Yip et al., 2019) and sustained attention (Rosenberg et al., 2016). Previous studies (e.g., (Rosenberg et al., 2018)) have used brain connectivity to develop predictive models of sustained attention that can be generalized to healthy and clinical populations. However, while behavioural changes in sustained attention have been documented and functional brain networks that predict substance use have been identified (Yip et al., 2019), the underlying change in sustained attention brain networks from adolescence to adulthood and their relation to substance use are relatively less well described. Lower sustained attention has been accompanied by both stronger reductions in neural activity in the visual cortex, as well as stronger recruitment of the right supramarginal gyrus with increasing time on a sustained attention task with central cues in cigarette smokers as opposed to non-smokers (Vossel et al., 2011). In a resting-state functional magnetic resonance imaging (fMRI) paradigm, cannabis users aged 16-26 had stronger connectivity between the left posterior cingulate cortex and the cerebellum, correlated with poorer performance on sustained attention/working memory and verbal learning measures (Ritchay et al., 2021). Although most brain connectomic research has utilized resting-state fMRI data, functional connectivity (FC) during task performance has demonstrated superiority in predicting individual behaviours and traits, due to its potential to capture more behaviourally relevant information (Dhamala et al., 2022; Greene et al., 2018; Yoo et al., 2018). Specifically, Zhao et al. (2023) suggested that task-related FC outperforms both typical task-based and resting-state FC in predicting individual differences. Hence, we applied task-related FC to predict sustained attention over time.

Previous studies found that FC patterns predicted individual differences in sustained attention (Chen et al., 2022; O’Halloran et al., 2018; Sripada et al., 2020), yet relatively little is known about the relationship between brain activity related to sustained attention and substance use over time. A latent change score model can quantify bidirectional longitudinal relations between substance use and both behaviours and brain activity associated with sustained attention, shedding light on how substance use impacts sustained attention and its associated brain activity, and vice versa. In this study, we used task-fMRI from the IMAGEN dataset, a longitudinal study with >1,000 participants at each timepoint (ages 14, 19, and 23 years-old). We first obtained task-related whole-brain connectivity and then used connectome-based predictive modeling (CPM) to predict sustained attention from ages 14 to 23. Additionally, previous cross-sectional and longitudinal studies (Broyd et al., 2016; Harakeh et al., 2012; Lisdahl and Price, 2012) have shown that there are linear relationships between substance use and sustained attention over time. We therefore employed correlation analyses and a latent change score model to estimate the relationship between substance use and both behaviours and brain activity associated with sustained attention. Given the substantial sample size and longitudinal design of Harakeh et al.’s study, we hypothesized that behavioural and predictive networks associated with lower sustained attention would predict increased substance use (particularly cigarette smoking) over time.

## Results

### 1. Behavioural changes over time

Reaction time (RT) variability is a straightforward measure of sustained attention, with increasing variability thought to reflect poor sustained attention. RT variability can be defined as the intra-individual coefficient of variation (ICV), calculated as the standard deviation of mean Go RT divided by the mean Go RT from Go trials in the stop signal task. Lower ICV indicates better sustained attention. Participants’ demographic information for all analyses is shown in Table 1 (see also Tables S1-S2). A linear mixed model analysis showed significant fixed effects of age (i.e., timepoint) on ICV (F_1895.3_ = 51.14, *P <* 0.001) (Fig. 1A). Post-hoc analysis showed that ICV decreased with age: ICV at age 14 was significantly higher than ICV at ages 19 (t = 6.535, *P <* 0.001) and 23 (t = 10.109, *P <* 0.001). ICV at age 19 was also significantly higher than that at age 23 (t = 4.768, *P <* 0.001). The full results of the linear mixed model analysis are shown in Tables S3-S4. In addition, we found that individual differences in ICV were significantly correlated between the three timepoints (Fig. 1B and Table S5, all *P <* 2.8e^-7^).

**Table 1.**
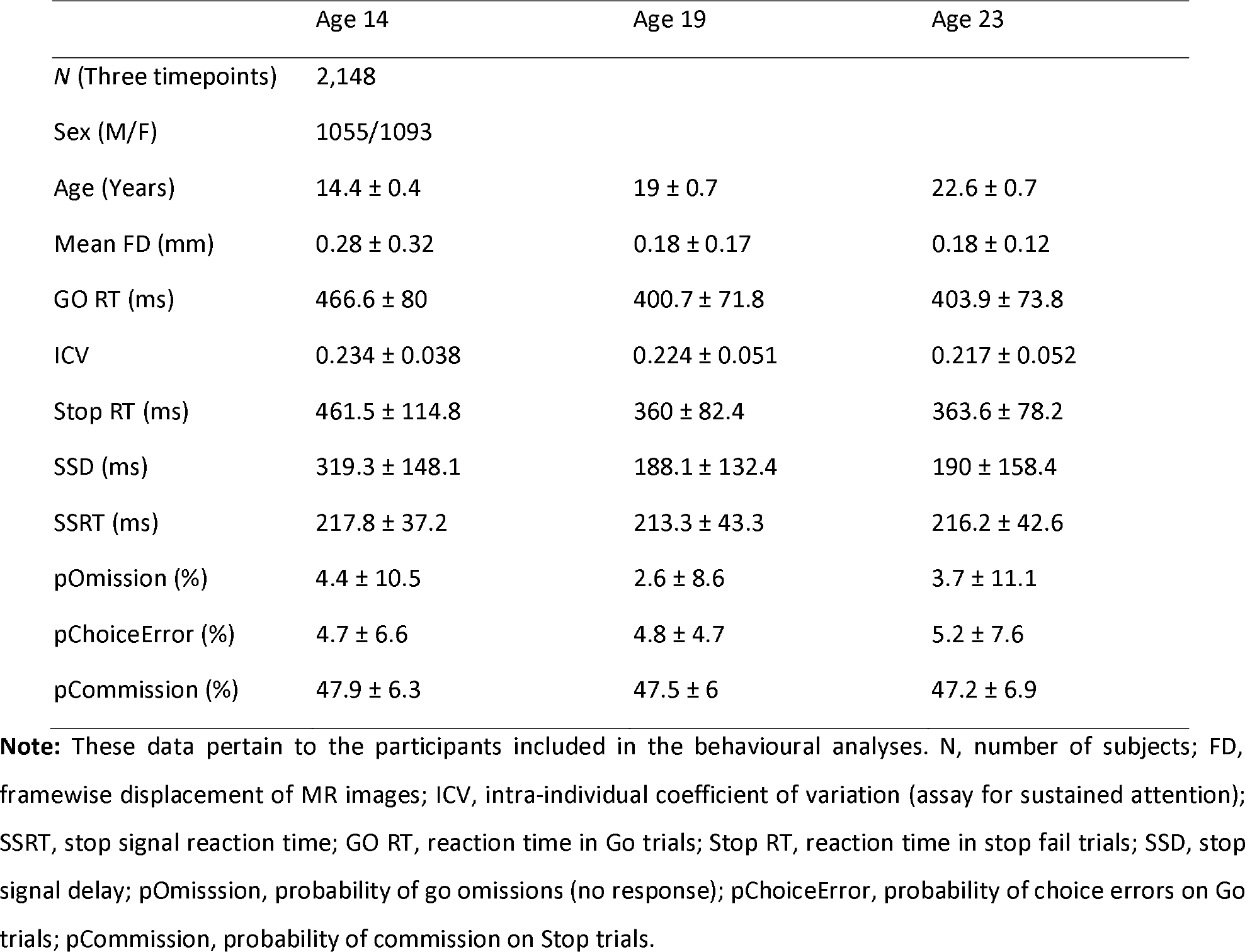
Demographic information of adolescents in the linear mixed model across three timepoints.

|  | Age 14 | Age 19 | Age 23 |
| --- | --- | --- | --- |
| *N* (Three timepoints) | 2,148 |  |  |
| Sex (M/F) | 1055/1093 |  |  |
| Age (Years) | 14.4 ± 0.4 | 19 ± 0.7 | 22.6 ± 0.7 |
| Mean FD (mm) | 0.28 ± 0.32 | 0.18 ± 0.17 | 0.18 ± 0.12 |
| GO RT (ms) | 466.6 ± 80 | 400.7 ± 71.8 | 403.9 ± 73.8 |
| ICV | 0.234 ± 0.038 | 0.224 ± 0.051 | 0.217 ± 0.052 |
| Stop RT (ms) | 461.5 ± 114.8 | 360 ± 82.4 | 363.6 ± 78.2 |
| SSD (ms) | 319.3 ± 148.1 | 188.1 ± 132.4 | 190 ± 158.4 |
| SSRT (ms) | 217.8 ± 37.2 | 213.3 ± 43.3 | 216.2 ± 42.6 |
| pOmission (%) | 4.4 ± 10.5 | 2.6 ± 8.6 | 3.7 ± 11.1 |
| pChoiceError (%) | 4.7 ± 6.6 | 4.8 ± 4.7 | 5.2 ± 7.6 |
| pCommission (%) | 47.9 ± 6.3 | 47.5 ± 6 | 47.2 ± 6.9 |

**Note:** These data pertain to the participants included in the behavioural analyses. *N*, number of subjects; FD, framewise displacement of MR images; ICV, intra-individual coefficient of variation (assay for sustained attention); SSRT, stop signal reaction time; GO RT, reaction time in Go trials; Stop RT, reaction time in stop fail trials; SSD, stop signal delay; pOmission, probability of go omissions (no response); pChoiceError, probability of choice errors on Go trials; pCommission, probability of commission on Stop trials.

**Fig. 1.**
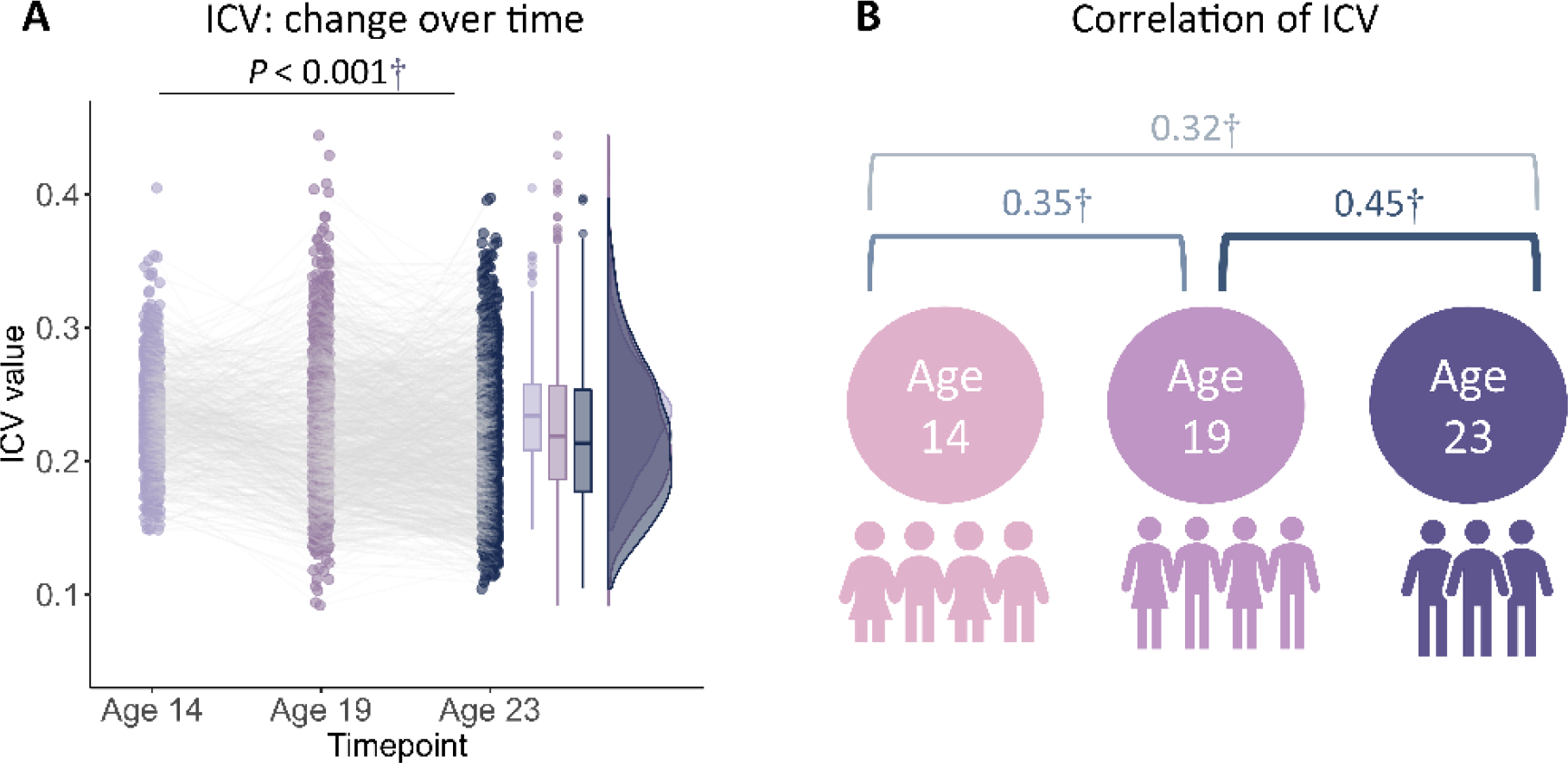
Intra-individual coefficient of variation (ICV) changes over time. (A) ICV changes over time. (B) Correlation of ICV between timepoints within subjects. †, P < 0.001.

### 2. Cross-sectional brain connectivity

This study employed CPM, a data-driven neuroscience approach, to identify three predictive networks— positive, negative, and combined — to predict ICV from brain functional connectivity. CPM typically uses the strength of the predictive networks to predict individual differences in traits and behaviors. The predictive networks were obtained based on connectivity analyses of the whole brain. Specifically, we assessed whether connections between brain areas (i.e., edges) in a task-related functional connectivity matrix derived from generalized psychophysiological interaction analysis were positively or negatively correlated with ICV using a significance threshold of *P* < 0.01. These positively or negatively correlated connections were regarded as positive or negative network, respectively. The network strength of positive networks (or negative networks) was determined for each individual by summing the connection strength of each positively (or negatively) correlated edge. The combined network was determined by subtracting the strength of the negative network from the positive network. We then built a linear model between network strength and ICV in the training set and applied these predictive networks to yield network strength and a linear model in the test set to calculate predicted ICV using k-fold cross validation.

Positive, negative, and combined networks derived from Go trials significantly predicted ICV: at age 14 (r = 0.25, r = 0.25, and r = 0.28, respectively, all *P <* 0.001) (Fig. 2A), at age 19 (r = 0.27, r = 0.25, r = 0.28, respectively, all *P <* 0.001) (Fig. 2B) and at age 23 (r = 0.38, r = 0.33, and r = 0.37, respectively, all *P <* 0.001) (Fig. 2C). The connectome patterns of predictive networks are shown in Figs. 2D-I. Fig. S2 summarizes the connectivity within and between functional networks and depicts their respective contribution to the predictive network. The above results were validated using 10-fold cross validation (CV); similar results were obtained when using 5-fold CV and leave-site-out CV (Table S6). The predictive networks had similar connectome patterns when different exclusion criteria for head motion were used (mean framewise displacement, mean FD < 0.2 −0.4 mm) (Figs. S4-S6A). In addition, we found that network strength of positive, negative, and combined networks derived from Go trials was significantly correlated between the three timepoints (Table S7, all *P <* 0.003).

**Fig. 2.**
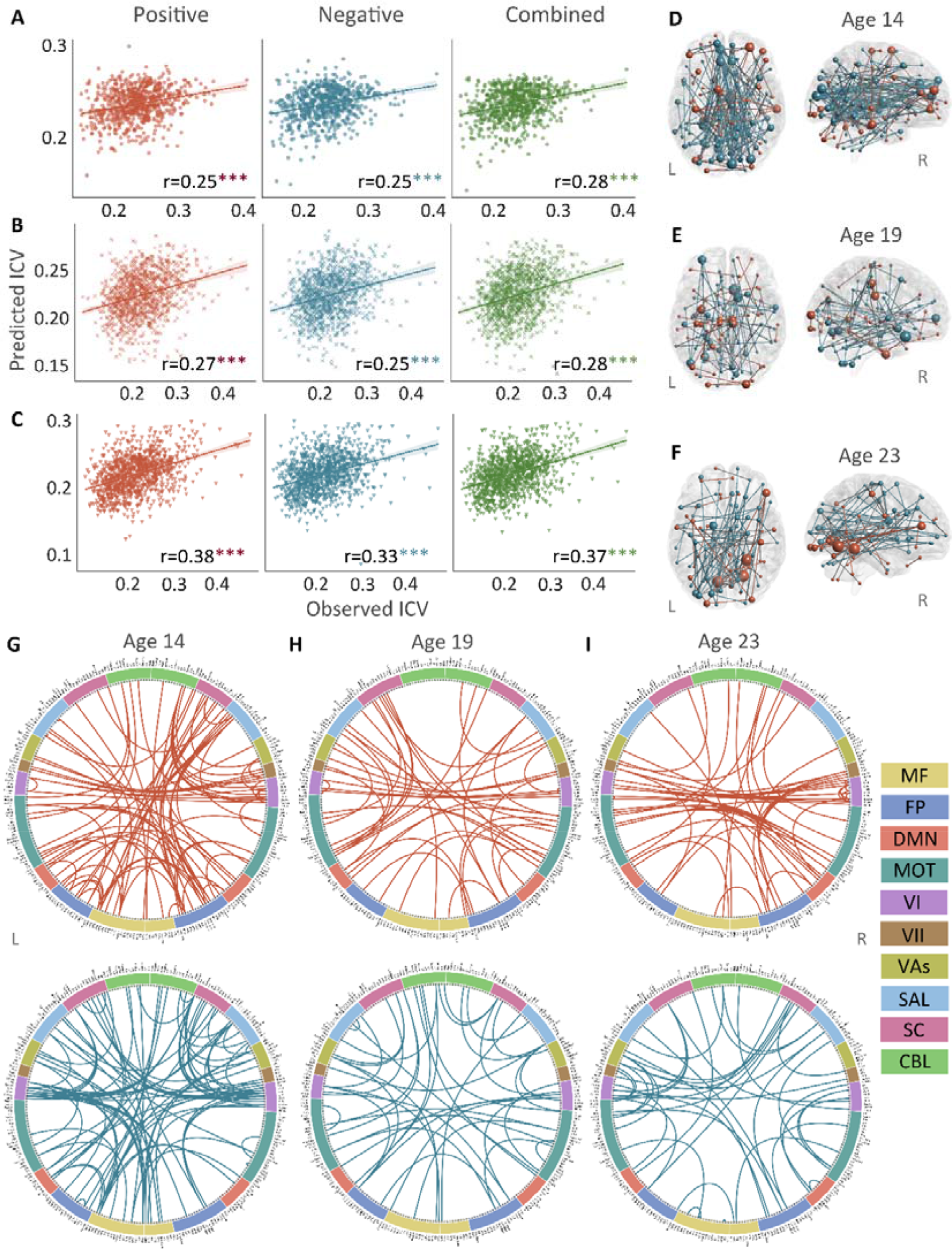
The predictive performances and networks of intra-individual coefficient of variation (ICV) per timepoint derived from Go trials. Correlation between observed and predicted ICV in positive, negative, and combined networks at (Panel A) age 14, (Panel B) age 19, and (Panel C) age 23. Predictive networks for ICV are at (Panel D) age 14, (Panel E) age 19, and (Panel F) age 23. Connectome of positive and negative networks of ICV at (Panel G) age 14, (Panel H) age 19, and (Panel I) age 23. The edges depicted above are those selected in at least 95% of cross-validation folds. Red, blue, and green spheres/lines/scatters represent positive, negative, and combined networks separately. MF, Medial frontal; FP, Frontoparietal; DMN, Default mode; MOT, Motor; VI, Visual I; VII, Visual II; VAs, Visual association; SAL, Salience; SC, Subcortical; CBL, Cerebellar. R/L, right/left hemisphere. ***, P < 0.001.

Positive, negative, and combined networks derived from Successful stop trials significantly predicted ICV: at age 14 (r = 0.22, *P <* 0.001; r = 0.12, *P =* 0.017; and r = 0.20, *P <* 0.001, respectively) (Fig. 3A), at age 19 (r = 0.19, *P <* 0.001; r = 0.15, *P =* 0.001; and r = 0.18, *P <* 0.001, respectively) (Fig. 3B), and at age 23 (r = 0.24, r = 0.21, and r = 0.23, respectively, all *P <* 0.001) (Fig. 3C). The connectome patterns of predictive networks are shown in Figs. 3D-I. Fig. S3 summarizes the connectivity within and between functional networks and the proportion of brain networks involved in the predictive network. We obtained similar results using a 5-fold CV and leave-site-out CV (Table S6). The predictive networks had similar connectome patterns when different exclusion criteria for head motion were used (mean FD < 0.2 −0.4 mm) (Figs. S4-S6B). In addition, we found that network strength of positive, negative, and combined networks derived from Successful stop trials was significantly correlated between the three timepoints (Table S7, all *P <* 0.001).

**Fig. 3.**
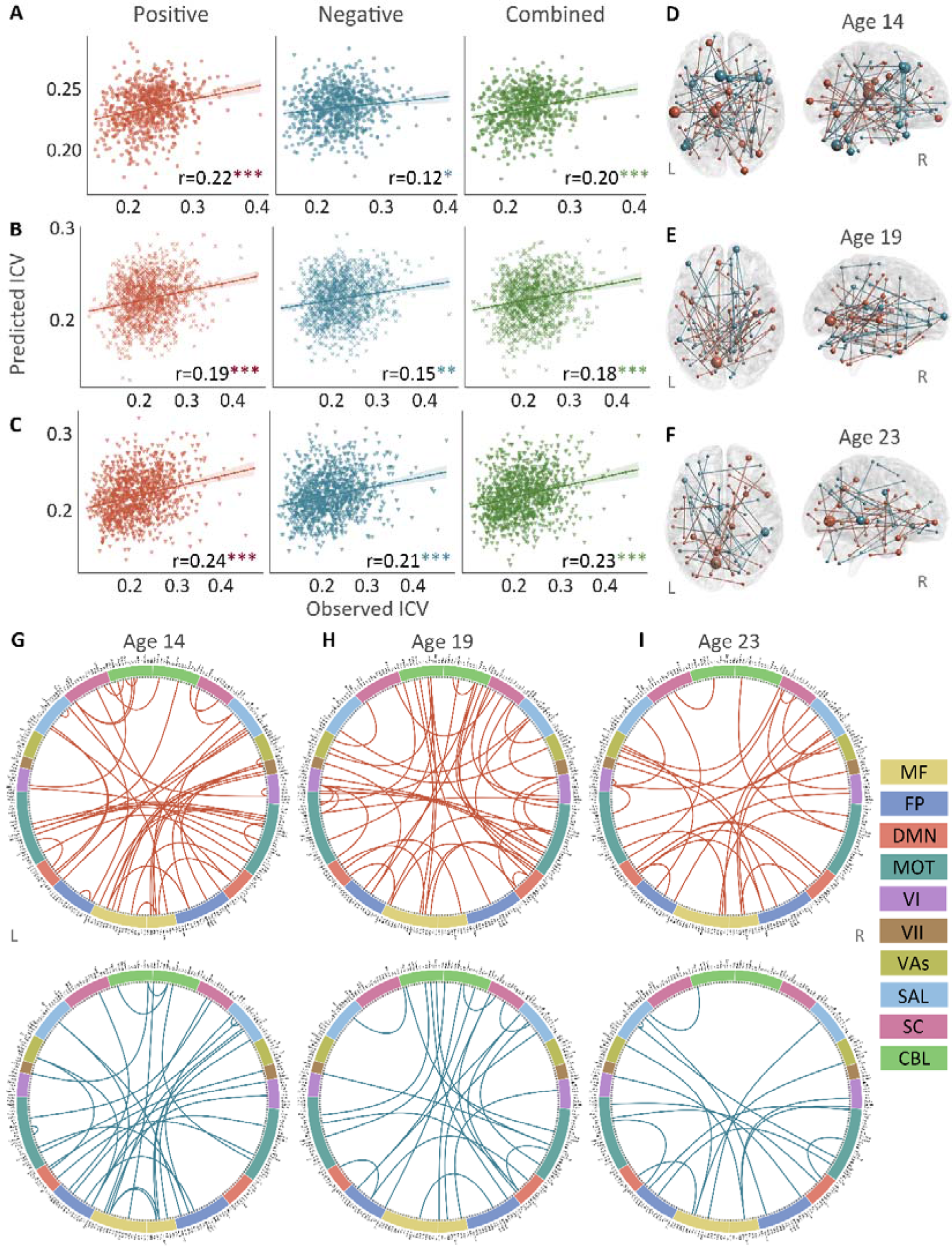
The predictive performances and networks of intra-individual coefficient of variation (ICV) per timepoint derived from Successful stop trials. Correlation between observed and predicted ICV in positive, negative, and combined networks at (Panel A) age 14, (Panel B) age 19, and (Panel C) age 23. Predictive networks for ICV are at (Panel D) age 14, (Panel E) age 19, and (Panel F) age 23. Connectome of positive and negative networks of ICV at (Panel G) age 14, (Panel H) age 19, and (Panel I) age 23. The edges depicted above are those selected in at least 95% of cross-validation folds. Red, blue, and green spheres/lines/scatters represent positive, negative, and combined networks separately. MF, Medial frontal; FP, Frontoparietal; DMN, Default mode; MOT, Motor; VI, Visual I; VII, Visual II; VAs, Visual association; SAL, Salience; SC, Subcortical; CBL, Cerebellar. R/L, right/left hemisphere. *, P < 0.05; **, P < 0.01; ***, P < 0.001.

To examine the specificity of sustained attention networks identified from CPM analysis, the correlations between the network strength of positive and negative networks and performances from a neuropsychology battery (CANTAB) (Fray et al., 1996) were calculated at each timepoint separately. All positive and negative networks derived from Go and Successful stop trials were significantly correlated with a behavioural assay of sustained attention – the RVP task – at ages 14 and 19 (all *P* values < 0.028). Age 23 had no RVP task data in the IMAGEN study. There were sporadic significant correlations between constructs such as delay aversion/impulsivity and negative network strength, for example, but the most robust correlations were with the RVP. Detailed information is shown in Supplementary materials and Table S12.

### 3. ICV prediction across time

Positive, negative, and combined networks derived from Go trials defined at age 14 predicted ICV at ages 19 (r = 0.16, r = 0.14, and r = 0.16, all *P <* 0.001) (Fig. 4A, top row) and 23 (r = 0.20, r = 0.12, and r = 0.17, all *P <* 0.001) (Fig. 4A, middle row) respectively. Likewise, positive, negative, and combined networks derived from Go trials defined at age 19 predicted ICV at age 23 (r = 0.30, r = 0.26, and r = 0.31, respectively, all *P <* 0.001) (Fig. 4A, bottom row).

Positive, negative, and combined networks derived from Successful stop trials defined at age 14 predicted ICV at age 19 (r = 0.11, r = 0.12, and r = 0.13, all *P <* 0.001) (Fig. 4B, top row) and 23 (r = 0.14, r = 0.15, and r = 0.15, all *P <* 0.001) (Fig. 4B, middle row) respectively. Positive, negative, and combined networks derived from Successful stop trials defined at age 19 predicted ICV at age 23 (r = 0.17, r = 0.16, and r = 0.17, respectively, all *P <* 0.001) (Fig. 4B, bottom row).

**Fig. 4.**
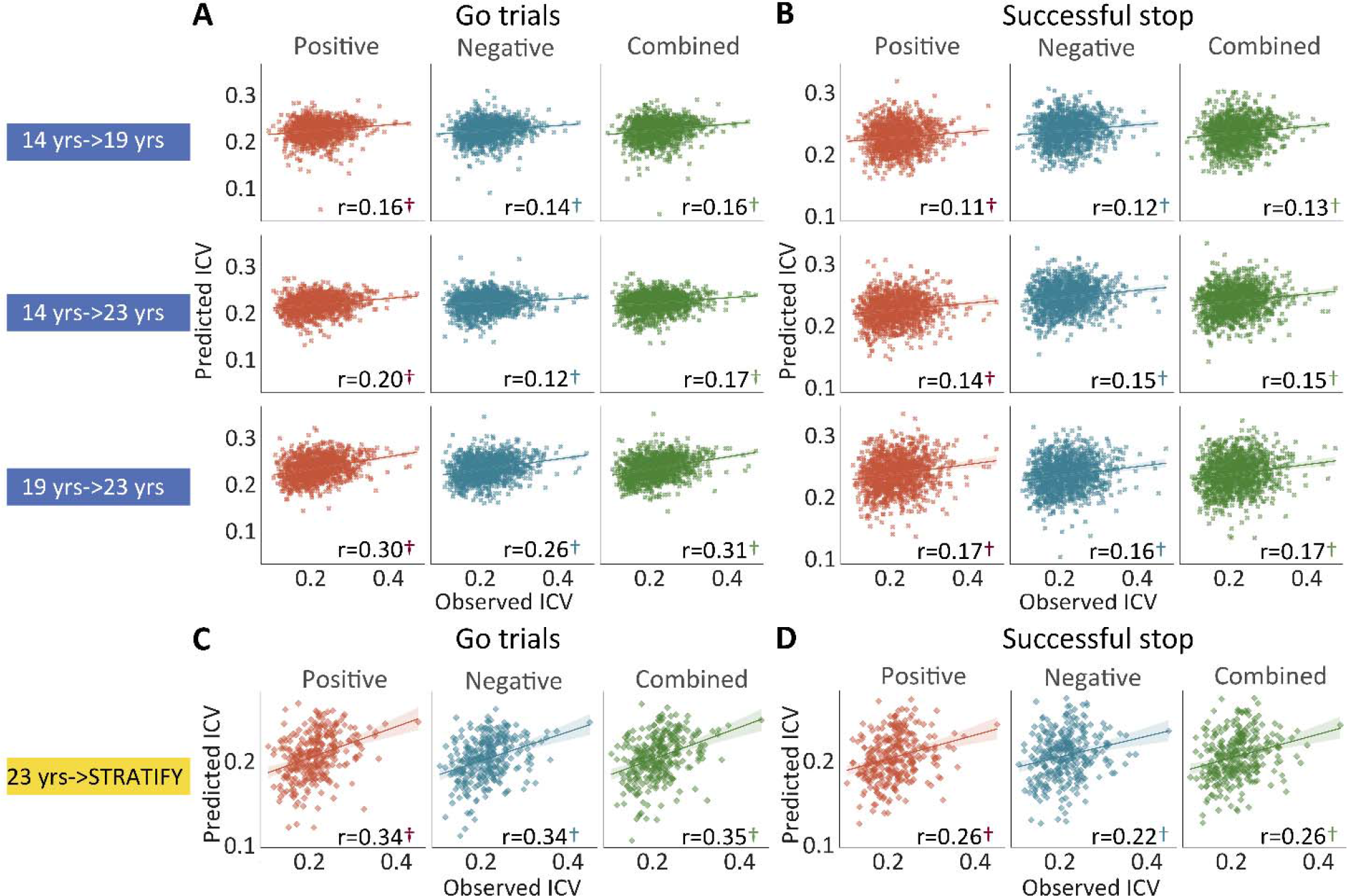
The predictive performances of intra-individual coefficient of variation (ICV) across timepoints and generalization in STRATIFY. Predictive performances of ICV (Panel A) derived from Go trials and (Panel B) derived from Successful stop trials. The top, middle, and bottom rows of (A) and (B) panels show the predictive performance: using models defined at age 14 to predict age 19 (i.e., 14 yrs -> 19 yrs), using models defined at age 14 to predict age 23 (i.e., 14 yrs -> 23 yrs), and using models defined at age 19 to predict age 23 (i.e., 19 yrs ->23 yrs) respectively. Generalization of predictive networks predicting ICV defined at age 23 in STRATIFY (i.e., 23 yrs -> STRATIFY) derived from (Panel C) Go trials and (Panel D) Successful stop trials. The red, blue, and green scatter represent positive, negative, and combined networks. †, P < 0.001.

### 4. Generalization of ICV brain networks

We tested if the predictive networks defined at age 23 in IMAGEN would generalize to an external dataset, namely STRATIFY (N = ∼300), comprising individuals also aged 23. When applied to the whole STRATIFY sample, positive, negative, and combined networks derived from Go trials at age 23 in IMAGEN predicted ICV in STRATIFY (r = 0.34, r = 0.34, and r = 0.35, respectively, all *P <* 0.001) (Fig. 4C), as did networks derived from Successful stop trials (r = 0.26, r = 0.22, and r = 0.26, respectively, all *P <* 0.001) (Fig. 4D).

### 5. Factor analysis of substance use

Exploratory factor analysis on data from the Timeline Followback (TLFB) (Sobell et al., 1996), an instrument for measuring the consumption of alcohol, drugs, and smoking for participants, yielded two common factors at age 14 and three common factors at ages 19 and 23. According to the rotated factor loading analysis, at age 14, two common factors were identified, which we labeled as (i) *alcohol* and (ii) *cigarette and cannabis* use (*Cig+CB)*. At ages 19 and 23, three common factors were identified, which we labeled as (i) *alcohol*, (ii) *Cig+CB*, and (iii) *drug* (including cocaine, ecstasy, and ketamine) use. Additional details about this data reduction step are shown in Fig. S7 and Table S11.

### 6. Correlation between behaviour and brain to cannabis and cigarette use

We calculated the Spearman correlation between ICV/sustained brain activity and TLFB factor score per timepoint and across timepoints. Brain activity was measured by the strength of positive and negative networks predicting sustained attention. The *P* values were corrected by false-discovery rate (FDR) correction (q < 0.05). Figs. 5A-C summarizes the results showing the correlation between ICV/brain activity and Cig+CB per timepoint and across timepoints. Fig. 5A shows correlations between ICV and Cig+CB (Tables S14-15). ICV was correlated with Cig+CB at ages 19 (Rho = 0.13, *P* < 0.001) and 23 (Rho = 0.17, *P* < 0.001). ICV at ages 14 (Rho = 0.13, *P* = 0.007) and 19 (Rho = 0.13, *P* = 0.0003) were correlated with Cig+CB at age 23. Cig+CB at age 19 was correlated with ICV at age 23 (Rho = 0.13, *P* = 9.38E-05). Fig. 5B shows correlations between brain activity derived from Go trials and Cig+CB (Tables S18-19). Brain activities of positive and negative networks derived from Go trials were correlated with Cig+CB at age 23 (positive network: Rho_p_ = 0.12, *P* < 0.001; negative network: Rho_n_ = −0.11, *P* < 0.001). Brain activity of the negative network derived from Go trials at age 14 was correlated with Cig+CB at age 23 (Rho_n_ = −0.16, *P* = 0.001). Cig+CB at age 19 was correlated with brain activity of the positive network derived from Go trials at age 23 (Rho_p_ = 0.10, *P* = 0.002). Fig. 5C shows the correlations between brain activity derived from Successful stop and Cig+CB (Tables S18-19). Brain activities of positive and negative networks derived from Successful stop were correlated with Cig+CB at ages 19 (positive network: Rho_p_ = 0.10, *P* = 0.001; negative network: Rho_n_ = −0.08, *P* = 0.013) and 23 (positive network: Rho_p_ = 0.13, *P* < 0.001; negative network: Rho_n_ = −0.11, *P* = 0.001). No correlation between alcohol use and ICV/brain activity was found after FDR correction. Detailed results on the correlation between ICV/brain activity and substance use can be found in the supplementary materials (Tables S14-21).

**Fig. 5.**
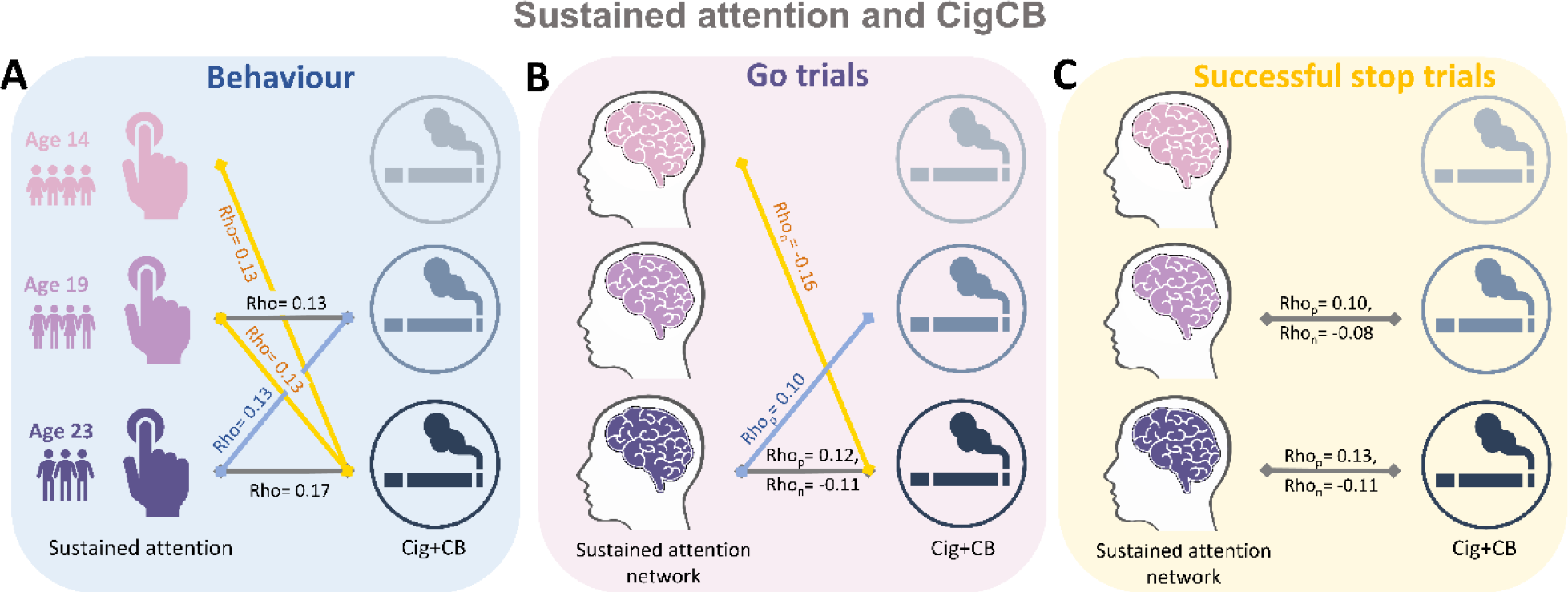
Significant correlations between sustained attention and substance use across timepoints (FDR correction, q<0.05). (A) Correlations between the intra-individual coefficient of variation (ICV) and cigarette and cannabis use (Cig+CB) across timepoints. Correlations between sustained attention network strength and Cig+CB across timepoints (B) derived from Go trials and (C) derived from Successful stop trials. Rho_p_: r value between network strength of the positive network. Rho_n_: r value between network strength of the negative network.

### 7. Bivariate latent change score model

We used a bivariate latent change score model to explore the relationship between substance use (specifically Cig+CB and alcohol use) and ICV/brain activity. This approach tests for bi-directional associations, examining how substance use at age 14 predicts changes in ICV/brain activity from ages 14 to 23 and vice versa (Fig. 6). Below, we present the findings regarding the lagged effects of substance use on ICV/brain activity and the lagged effects of ICV/brain activity on substance use (Table 2). The *P* values were corrected by FDR correction (q < 0.05).

**Fig. 6.**
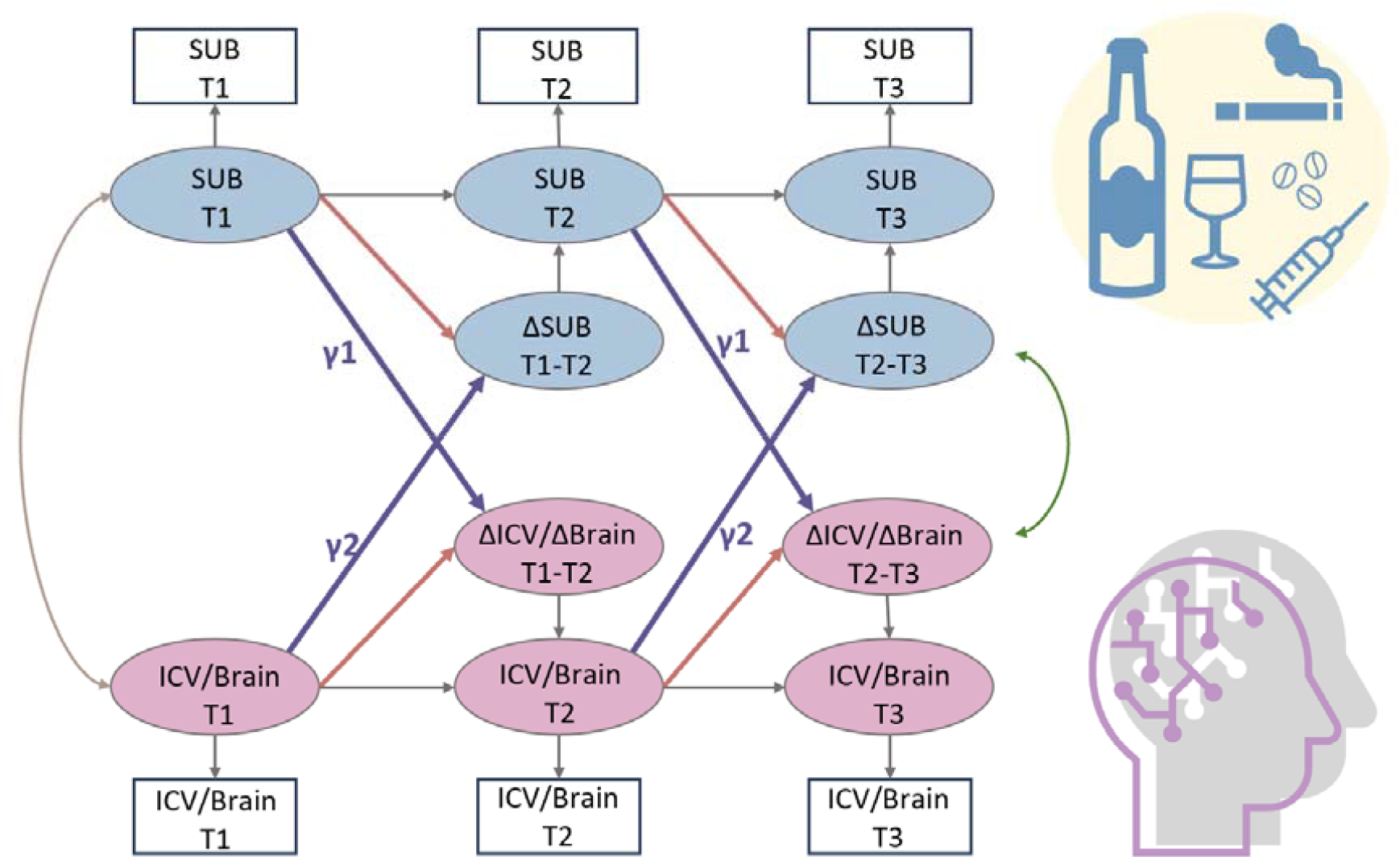
A simplified bivariate latent change score model for substance use and ICV/brain activity. SUB, substance use (alcohol, cigarette and cannabis use); Brain, brain network strength of positive/negative network of sustained attention derived from Go trials/Successful stop trials. ICV, intra-individual coefficient of variation. T1, Timepoint 1 (age 14); T2, Timepoint 2 (age 19); T3, Timepoint 3 (age 23). γ1, lagged effects of substance use on ICV or brain activity. γ2, lagged effects of ICV or brain activity on substance use. The square/circle represents the observation/true score in the model.

#### 7.1 Lagged effects of Cig+CB on changes in ICV and brain activity

We examined if Cig+CB use at age 14 predicted the changes in ICV or brain activity (i.e., predictive network strength) associated with sustained attention across ages 14-23. No significance was observed in the lagged effects of Cig+CB on changes in ICV and brain activity (all *P* > 0.172).

#### 7.2 Lagged effects of ICV and brain activity on changes in Cig+CB

We examined if ICV or brain activity associated with sustained attention at age 14 predicted changes in Cig+CB use across ages 14-23. Behaviours and brain activity associated with poor sustained attention predicted a greater increase in subsequent cigarette and cannabis use. Specifically, higher ICV at age 14 predicted a greater increase in Cig+CB from ages 14 to 23 (*Std. β* = 0.12, *P* < 0.001). Higher sustained attention network strength for positive network derived from Go trials at age 14 predicted a greater increase in Cig+CB from ages 14 to 23 (*Std. β* = 0.09, *P* = 0.006). Lower sustained attention network strength for the negative network, also derived from Go trials at age 14, predicted a greater increase in Cig+CB from ages 14 to 23 (*Std. β* = −0.09, *P* = 0.006). No other lagged effects of brain activity on changes in Cig+CB remained significant after FDR correction (all *P* > 0.047). Fig. 7 illustrates the changes in raw scores of cigarette and cannabis use from the TLFB for individuals at age 14 with higher sustained attention (i.e., lower ICV, lower strength of positive network, or higher strength of negative network) and lower sustained attention (i.e., higher ICV, higher strength of positive network, or lower strength of negative network).

**Fig. 7.**
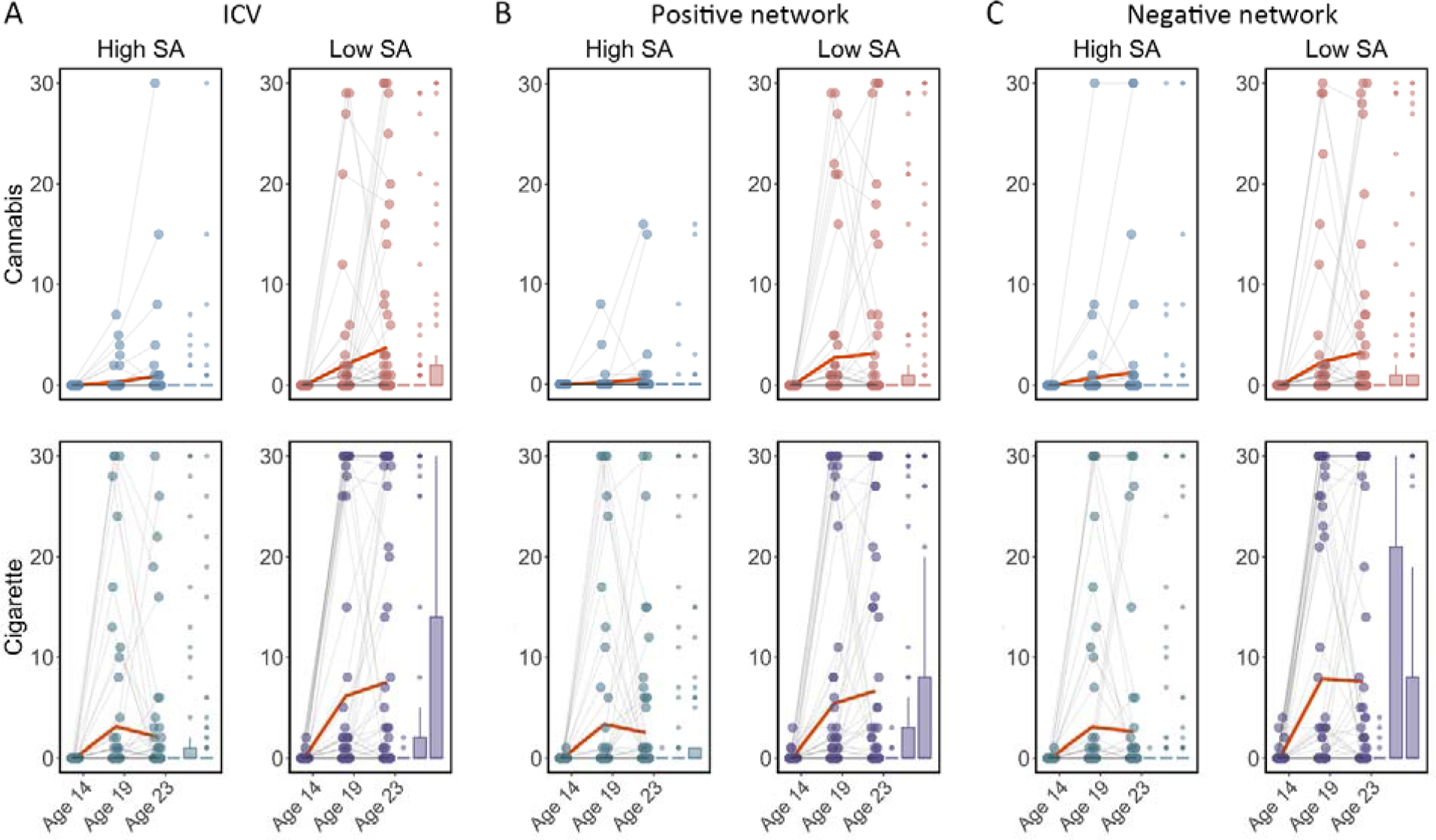
Cigarette and cannabis score in Timeline Followback changes in individuals with high sustained attention (High SA) and low sustained attention (Low SA) from ages 14 to 23. Participants were categorized into five equal groups based on the intra-individual coefficient of variation (ICV), strength of positive network, and strength of negative network at age 14. (A) Top ICV (Low SA) and bottom ICV (High SA) groups. (B) The top strength of the positive network (Low SA) and bottom strength of the positive network (High SA) groups derived from Go trials. (C) The top strength of the negative network (High SA) and bottom strength of the negative network (Low SA) groups derived from Go trials. Note that the higher strength of the negative network reflects lower ICV and higher sustained attention.

#### 7.3 Association between alcohol use and ICV/brain activity

We examined if alcohol use at age 14 predicted changes in ICV or brain activity associated with sustained attention across ages 14-23, or vice versa. No significant results were found for the lagged effects of alcohol use on changes in ICV and brain activity, nor the lagged effects of ICV and brain activity on changes in alcohol use. The *P* values were insignificant after FDR correction (all *P* > 0.011).

**Table 2.**
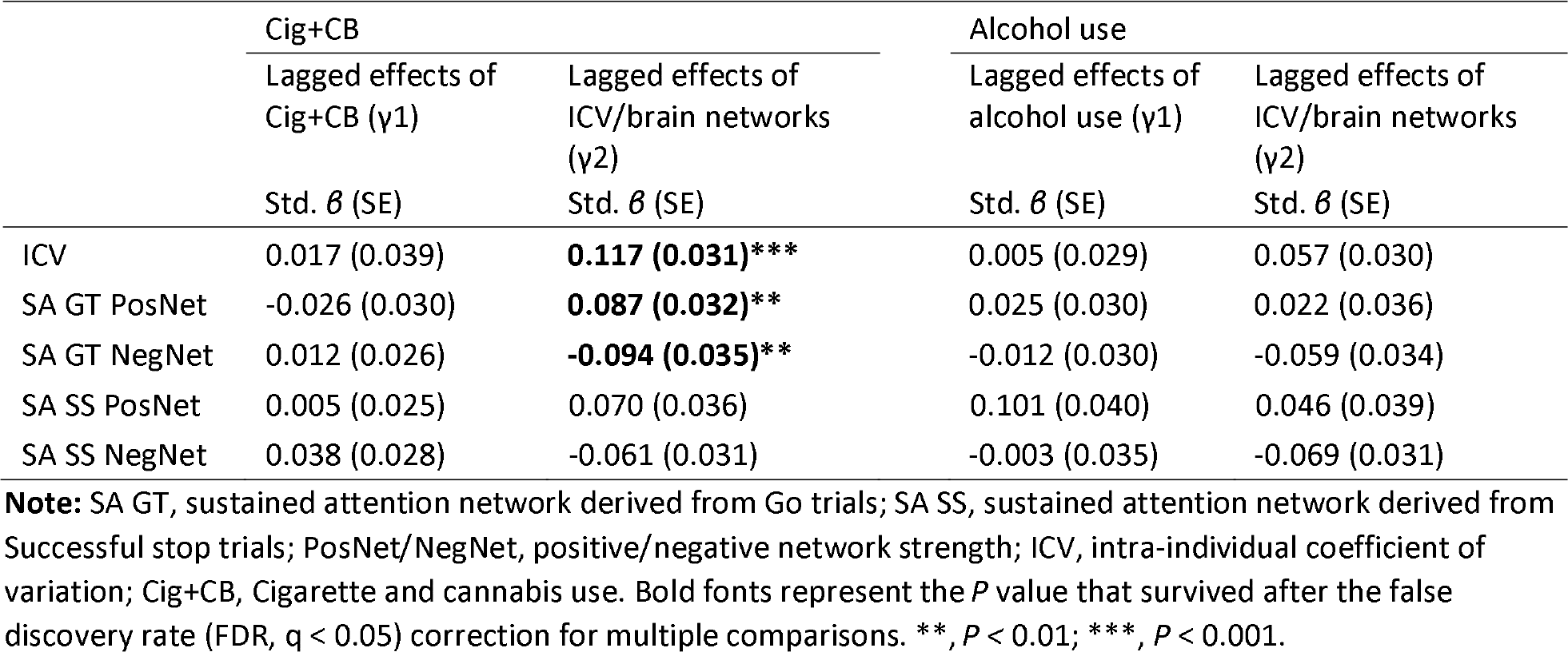
Bivariate latent change score model showing the bi-directional association between substance use and ICV/brain networks (False Discovery Rate corrected)

## Discussion

It is well known that increased substance use, including cigarettes and cannabis, is associated with poorer sustained attention in late adolescence and early adulthood (Chamberlain et al., 2012; Dougherty et al., 2013). However, previous studies, which were predominantly cross-sectional or under-powered, left a critical question unanswered. That is, was the impairment in sustained attention a predictor of substance-use or a marker of the inclination to engage in such behaviour? Using a substantial sample size, our results indicate that behaviour and brain connectivity associated with poorer sustained attention at age 14 predicted a larger increase in cannabis and cigarette smoking from ages 14-23. Furthermore, our findings highlight the robustness of the brain network associated with sustained attention over time, making the latter a potentially useful biomarker for vulnerability to substance use.

### Substance use and the sustained attention network

Our study applied a latent change score model on a large longitudinal dataset, testing the precedence between substance use and sustained attention. In contrast to prior research suggesting that substance use impaired sustained attention (Broyd et al., 2016; Figueiredo et al., 2020), our results indicate that lower sustained attention also predates substance use. A link between substance use and sustained attention is plausible, given the underlying neurobiology of this sustained attention. Substantial evidence from neuropharmacological studies in rats and humans has shown the modulatory role of neurotransmitters in sustained attention (Bloomfield et al., 2016; Granon et al., 2000; Marshall et al., 2019). Elevated dopamine and noradrenaline levels in the prefrontal cortex lead to improved sustained attention in a dose-dependent manner (Marshall et al., 2019). In humans, methylphenidate, a psychostimulant commonly used to treat ADHD, increases both noradrenaline and dopamine signalling and improves sustained attention (Dockree et al., 2017). Thus, poorer sustained attention may reflect a lower basal level of dopamine and noradrenaline. More importantly, studies in primates (Morgan et al., 2002; Nader et al., 2006), rodents (Dalley et al., 2007; Trifilieff et al., 2017), and humans (Casey et al., 2014; Trifilieff and Martinez, 2014; Volkow et al., 2006) have indicated that low basal dopamine levels are markers of vulnerability for increased drug administration. For example, Casey et al. (2014) demonstrated that blunted dopamine release may precede the development of addiction in humans. Nader et al. (2006) found a negative correlation between baseline D2 receptor availability and rates of cocaine self-administration in monkeys. Thus, these findings collectively suggest that sustained attention and its brain network could serve as a biomarker of vulnerability to substance use.

These results emphasize the specificity of sustained attention and its associated brain networks, rather than other cognitive abilities, for predicting substance use over time. Unlike sustained attention, no significant differences in cigarette and cannabis use were observed between individuals with lower and higher working memory at baseline during the Strategy working memory task (Table S23 and Fig. S8). Our results support the behavioural-only findings of a previous study (Harakeh et al., 2012), which found that individuals with poorer sustained attention, rather than other cognitive functions, were more likely to initiate smoking cigarettes. Our study goes further by showing that sustained attention brain networks can predict substance use in the future.

### Neural associations between cigarette and cannabis use

We constructed composite scores of substance use. An exploratory factor analysis identified cigarettes and cannabis items as a common factor, aligning with previous studies (Ferland and Hurd, 2020; Hindocha et al., 2016; Weinberger et al., 2018) that indicate concurrent cannabis and cigarette use among users. A national survey in America indicated that 18–23% of cigarette smokers aged 12–17 met the criteria for cannabis use disorder, in contrast to only 2% of non-smoking youth (Weinberger et al., 2018). Another national online survey in the UK reported that 80.8% of cigarette smokers engage in cannabis consumption, indicating a prevalent practice of co-administering cannabis and tobacco through smoking (Hindocha et al., 2021). Shared genetic factors (Agrawal et al., 2010; Stringer et al., 2016) and similar neural associations (Wetherill et al., 2015) contribute to the co-use of cannabis and cigarettes. Stringer et al. (2016) demonstrated a strong and significant genetic correlation between lifetime cannabis use and lifetime cigarette smoking within a large cohort of 32,330 subjects, suggesting a high degree of genetic sharing between the two. Using neuroimaging techniques, Wetherill et al. (2015) indicated that individuals who used cannabis, smoked tobacco, or engaged in co-use exhibited larger gray matter volumes in the left putamen compared to healthy controls. Both nicotine and cannabis have similar effects on mesolimbic dopaminergic pathways engaged, modulating dopamine release in the striatum (Bossong et al., 2009; Dongelmans et al., 2021). Collectively, these findings suggest a similar neural association between cigarette and cannabis use.

### Specificity and robustness of sustained attention networks

The brain networks we describe were specific to sustained attention. The strength of the sustained attention brain network was robustly correlated with RVP task performance, a typical sustained attention task, rather than other cognitive measures (Table S12). Importantly, as highlighted in a previous study (Cwiek et al., 2022), emphasizing the importance of generalization in an external dataset, our study found that the sustained attention network derived from Go trials and Successful stop trials generalized to an external dataset (See further discussion on the generalization in subgroups in STRATIFY in Supplemental materials).

We also replicated and extended the developmental pattern of sustained attention and its networks from mid-adolescence to young adulthood. A notable enhancement in sustained attention (i.e., decreased ICV) was observed from ages 14 to 23, as expected (Fortenbaugh et al., 2015; Williams et al., 2005). Sustained attention networks derived from Go and Successful stop trials predicted behaviour at different timepoints, implying that individual differences in sustained attention and associated networks were preserved throughout development. Previously, in neurodiverse youth, attention networks in individuals remained stable across months to years (Horien et al., 2022). Rosenberg et al. (2020) also illustrated that the same functional connections predicting overall sustained attention ability also forecasted attentional changes observed over minutes, days, weeks, and months. Here, we contribute to these insights by extending the understanding that attention-network stability is not only applicable to neurodiverse populations but also holds in a sizeable cohort of healthy participants. Furthermore, our findings indicate that sustained attention networks remain stable over several years, providing valuable insights into the potential for sustained attention to function as a robust and efficient biomarker for substance use. However, there are still some individual variabilities not captured in this study, which could be attributed to the diversity in genetic, environmental, and developmental factors influencing sustained attention and substance use. Future research should aim to explore these variabilities in greater depth to gain better understanding of the relationship between sustained attention and substance use.

In conclusion, robust sustained attention networks were identifiable from ages 14 to 23. Individual differences in sustained attention network strength were predictable across time. Poorer sustained attention and strength of the associated brain networks at age 14 predicted greater increases in cannabis and cigarette smoking from ages 14 to 23.

## Materials and Methods

### 2.1 Participants

All neuroimaging data and behavioural data were obtained from the IMAGEN study. IMAGEN is a large longitudinal study that recruited over 2000 participants aged 14 to 23 in Europe (Kaiser et al., 2022). This study used the stop signal task fMRI data at ages 14, 19, and 23. In addition, we used an independent dataset STRATIFY as external validation for age 23. STRATIFY (N = ∼300) is a sub-dataset within IMAGEN that recruits fMRI data from patients aged 23. The IMAGEN study conformed to the ethical standard outlined by the Declaration of Helsinki and was approved by ethics committees at each site. Written informed consent was obtained from participants and their parents. We followed the exclusion criteria outlined in previous studies (O’Halloran et al., 2018; Whelan et al., 2014). Participants were excluded from the CPM analysis if they had more than 20% errors on the Go trials (incorrect responses or responses that were too late) or if they had a mean framewise displacement (mean FD) > 0.5 mm. Finally, 717 subjects at age 14, 1081 subjects at age 19, and 1120 subjects at age 23 were used to predict ICV. In STRATIFY, 304 subjects were used to predict ICV.

### 2.2 Stop signal task

The stop signal task required subjects to respond to a Go signal (arrows pointing left/right) by pressing the left/right button while withholding their response if the Go signal was unpredictably followed by a stop signal (arrows pointing upwards). The Go signal was displayed on the screen for 1000 ms in the Go trials, while the stop signal appeared for 100-300 ms following the go signal on average 300 ms later in unpredictable stop trials. To adjust task difficulty dynamically, we used a tracking algorithm on the delay between the Go signal and stop signal (stop signal delay, SSD, 250-900 ms in 50 ms increments) (Verbruggen et al., 2019), to produce 50% successful and 50% unsuccessful inhibition trials. The task at age 14 included 400 Go trials and 80 variable delay stop trials, with 3 and 7 Go trials between successive stop trials. The task at ages 19 and 23 consisted of 300 Go trials and 60 variable delay stop trials. Before the MRI scan, subjects also performed a practice session with a block of 60 trials to become familiar with the task. ICV is used to assess sustained attention in this task for each subject. ICV reflects short-term within-person variations in task performance (Verbruggen et al., 2019). Specifically, ICV is computed by dividing the standard deviation of mean go RT by the mean go RT. Lower ICV indicates better sustained attention.

### 2.3 Self-Report Questionnaires

#### 2.3.1 Puberty development scale (PDS)

The PDS, an 8-item self-report assessment, measures the pubertal development of adolescents (Petersen et al., 1988). The PDS evaluates physical development using a 5-point scale where 1 corresponds to prepubertal, 2 to beginning pubertal, 3 to mid-pubertal, 4 to advanced pubertal, and 5 to post pubertal. In addition, the items are adapted for sex, such as voice changes for males or menarche for females.

#### 2.3.2 Timeline Followback (TLFB)

We used the TLFB, a retrospective self-report instrument that uses a calendar method to evaluate prior substance use consumption over the past 30 days (Sobell et al., 1996). The TLFB has strong reliability and validity for assessing alcohol consumption, and we used it to measure the use of alcohol, drugs, and smoking for subjects.

### 2.4 MRI acquisition and pre-processing

Functional MRI data of the stop-signal task in the IMAGEN study were collected at eight scan sites (London, Nottingham, Dublin, Mannheim, Dresden, Berlin, Hamburg, and Paris), and data in STRATIFY were collected at three scan sites (Berlin, two scanners in London) with 3T MRI scanners. The MR scanning protocols, cross-site standardization, and quality checks are further described in (Whelan et al., 2012). All images were obtained using echo-planar imaging (EPI) sequence with the following parameters: repetition time (TR) = 2.2s, echo time (TE) = 30ms, flip angle = 75°, field of view (FOV) = 224 mm × 224 mm, data matrix = 64 × 64, slice thickness = 2.4 mm with 1 mm slice gap, voxel size = 3.5 × 3.5 × 4.38 mm, 40 transversal interleaved slices. The MRI data has 444 volumes at age 14 and 320-350 volumes at ages 19 and 23. Standardized hardware was used for visual stimulus presentation (Nordic Neurolab, Bergen, Norway) at all scan sites.

All fMRI data from the IMAGEN study were pre-processed centrally using SPM12 (Statistical Parametric Mapping, http://www.fil.ion.ucl.ac.uk/spm/) with an automated pipeline. The images were corrected for slice timing and then realigned to the first volumes to correct head motions. Subjects were excluded from the study if they had a mean FD > 0.5mm. Subsequently, the data were non-linearly transformed to the Montreal Neurological Institute Coordinate System (MNI) space using a custom EPI template with the voxels resampled at 3 × 3 × 3 mm resolution. Finally, the images were smoothed with a Gaussian kernel at a full-width-at-half-maximum (FWHM) of 5 mm.

### 2.5 Generalized psychophysiological interaction (gPPI) analysis

In this study, we adopted gPPI analysis to generate task-related FC matrices and applied CPM analysis to investigate predictive brain networks from adolescents to young adults. PPI analysis describes task-dependent FC between brain regions, traditionally examining connectivity between a seed region of interest (ROI) and the voxels of the whole rest brain. However, this study conducted a *generalized* PPI analysis, which is on ROI-to-ROI basis (Di et al., 2021), to yield a gPPI matrix across the whole brain instead of just a single seed region. First, we conducted a general linear model (GLM) analysis on the pre-processed fMRI data to examine brain activity during the stop signal task. Two separate GLMs were created for Go trials and Successful stop trials. The Go trials model included three task regressors (Go trials, Failed stop trials, and Successful stop trials) and 36 nuisance regressors, which accounted for factors such as head motion and the signal from white matter and cerebrospinal fluid. The 36 nuisance regressors are 3 translations, 3 rotations, mean white matter signal, mean cerebrospinal fluid signal, mean grey matter signal, their derivatives, and the squares of all these variables. Given the high frequency of Go trials in SST, it is common to treat Go trials as an implicit baseline, as in previous IMAGEN studies (D’Alberto et al., 2018; Whelan et al., 2012). Hence, we built a separate GLM for Successful stop trials, which included two task regressors (Failed and Successful stop trials) and 36 nuisance regressors. All task regressors were modeled by convolving with the canonical hemodynamic response function (HRF) and high pass filtered (128 s). We then conducted a gPPI analysis across the entire brain using the Shen atlas with 268 regions (Shen et al., 2013) for both Go and Successful stop trials. The gPPI analysis involved deconvolving the time series of each ROI with the hemodynamic response function, multiplying it by the psychological variables of interest to yield a neural level PPI term, and convolving the resulting PPI term with the HRF to obtain the BOLD level PPI effects (Di and Biswal, 2019). Separate GLM models were used to estimate the PPI effect of each ROI for Go trials and Successful stop trials, regressing the eigenvariate of the seed ROI. The GLM of the Go trials included 1 regressor of another ROI eigenvariate, 3 regressors of task condition, 3 regressors of the PPI effects, and one contrast term (Equation 1). The GLM of Successful stop trials included 1 regressor of another ROI eigenvariate, 2 regressors of task condition, 2 regressors of the PPI effects, and one contrast term (Equation 2), shown as follows:

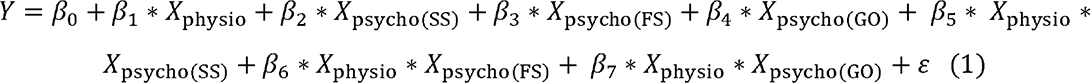

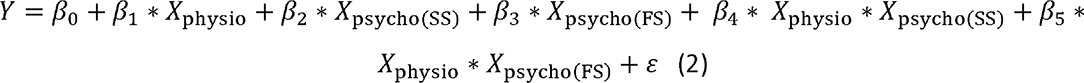

Note: SS, Successful stop trials; FS, Failed stop trials; GO, Go trials.

Where *γ* is the time series of seed ROI, *X_physio_* is the time series of another ROI, *X_physio_* is the task design term, and ε is the residual term. The generalized PPI analysis was performed across each ROI from the Shen atlas, resulting in a 268*268 PPI matrix for each subject derived from Go trials and Successful stop trials separately. The matrices were transposed and averaged with the original matrices to yield symmetrical matrices (Di et al., 2021), and prepared for further analysis.

### 2.6 Connectome-based predictive modeling

#### 2.6.1 ICV prediction

CPM is a data-driven method that can examine individual differences in brain connectivity (Shen et al., 2017). CPM identifies pairwise connections between brain regions most highly correlated with a given phenotype. Using the PPI matrix, we employed CPM to predict ICV, for ages 14, 19, and 23. The CPM analysis process includes feature selection, model building, and validation (Fig. S1). We applied cross-validation to divide all participants into training and testing sets. (i) First, we used partial correlation to calculate the relationship between each edge in the gPPI matrix and behavioural phenotype while controlling several covariates in the training set. These covariates included ages, genders, mode-centered PDS (at age 14 only), mean FD, and scan sites, regarded as a dummy variable. The r value with an associated *P* value for each edge was obtained, and a threshold *P* = 0.01 (Feng et al., 2024; Ren et al., 2021; Yoo et al., 2018) was set to select edges. The positive or negative correlated edges in feature selection were regarded as positive or negative networks. (ii) Second, we calculated network strength for each subject in the training set by summing the selected edges in the gPPI matrix for both positive and negative networks. We also estimated the network strength of a combined network by subtracting the strength of the negative from the strength of the positive network. (iii) Finally, we constructed predictive models based on the assumption of a linear relationship between network strength of the positive, negative, and combined networks, and behavioural phenotype in the training set. The covariates were also adjusted in this linear model. The network strengths for each subject in the testing set were calculated and input into the predictive model along with the covariates to predict each network’s behavioural phenotypes.

#### 2.6.2 Three cross-validation schemes

We used three CV schemes to test the robustness of predictive performance: k-fold (10-fold and 5-fold) and leave-site-out CV. For the k-fold CV, we randomly divided subjects into ten or five approximately equal-sized groups. For each fold, we trained the model on nine or four groups, respectively, and used it to predict the behavioural phenotype of the remaining group. We then assessed the predictive performance by comparing the predicted and observed values. For the leave-site-out CV, we divided subjects into eight groups based on their scan site. To account for the random splits of the k-fold CV, we repeated the process 50 times and calculated the average predictive performance for both the 10-fold and 5-fold CV (Lichenstein et al., 2021). In addition, we set a 95% threshold for selecting edges present in at least 48 out of 50 iterations to visualize the results. We also ran the CPM analysis with mean FD thresholds of 0.2, 0.3, and 0.4 mm to account for the influence of head motion on the predictive performance. Furthermore, we conducted the CPM analysis using a range of thresholds for feature selection and observed similar results across different thresholds (See supplementary materials Table S8). The main text shows the results of the 10-fold CPM. The 5-fold CPM and leave-site-out CV results are shown in supplementary materials.

#### 2.6.3 Prediction across timepoints and STRATIFY

To assess the ability of models developed at one timepoint to predict ICV at different timepoints, we applied predictive models developed at ages 14 and 19 to predict ICV at subsequent timepoints. Specifically, we used predictive models (including the parameters and selected edges) developed at age 14 to predict ICV at ages 19 and 23. We first calculated the network strength using the gPPI matrix at age 19 or 23 based on the selected edges identified from CPM analysis at age 14. We then used the linear model parameters (slope and intercept) from CPM analysis at age 14 to fit the network strength and predict ICV at age 19 or 23. Finally, we evaluated the predictive performance by calculating the correlation between the predicted and observed values at age 19 or 23. Similarly, we applied models developed at age 19 to predict ICV at age 23. In addition, we examined the generalizability of predictive models at age 23 by applying them to the STRATIFY dataset, which also includes subjects who were 23 years old. Furthermore, we estimated the predictive performances of ICV across patient groups in the STRATIFY. The correlation between the residual network strength of predictive networks and ICV was calculated across groups in the STRATIFY. The covariates, including age, sex, and mean FD, were regressed for network strength before the correlation analysis. It is worth noting that when applying models developed at one timepoint to predict at another timepoint or to generalize to a different dataset, the model was built using all subjects from the timepoint at which the model was developed.

### 2.7 Statistical analysis

#### 2.7.1 Exploratory Factor Analysis

To explore the underlying structure of adolescent substance use, we performed an exploratory factor analysis using principal component extraction (Gaskin and Happell, 2014) on TLFB using Predictive Analytics Software (SPSS) version 20. Factor analysis explores the underlying structure of a set of observed variables without imposing a preconceived structure on the outcome. We used six items at age 14 and nine items at ages 19 and 23 of TLFB, including alcohol, tobacco, cannabis, cocaine, ecstasy, and ketamine (as shown in Table S11). We excluded items assessing the use of other drugs due to high proportions of missing data, standard deviations close to 0, or a Kaiser-Meyer-Olkin (KMO) statistic for individual variables below 0.5, considered the minimum value for a sample to be adequate. The KMO measure of sampling adequacy was 0.66 at age 14, 0.81 at age 19, and 0.77 at age 23. In addition, all Bartlett’s tests of sphericity were significant (Age 14: χ^2^ (15) = 5137.067, *P* < 0.001; Age 19: χ^2^ (36) = 5031.641, *P* < 0.001; Age 23: χ^2^ (36) = 5106.265, *P* < 0.001), indicating that there was an underlying correlation structure, and that factor analysis was appropriate. We rotated the factors using the varimax method with kaiser normalization to make it easier to discern the underlying measured constructs.

#### 2.7.2 Linear mixed model

We constructed a linear mixed model to examine the change in ICV over time using the *lme4* and *lmerTest* packages in RStudio (version: 1.4; http://www.rstudio.com/) and R (version 4.1.1; https://www.r-project.org/). The timepoint was the fixed effect of interest in the model, while the subject was a random effect. Several covariates, including sex, scan sites, mode-center PDS, and age at 14, were also included as fixed effects in the models. The linear mixed model is shown as follows:

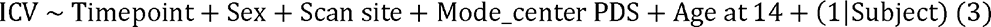

#### 2.7.3 Correlation between network strength and substance use

To examine the relationship between ICV/brain activity and substance use, we correlated the network strength of predictive networks with the factor scores of substance use at each timepoint and across all three timepoints separately. To control for potential confounders, we calculated residual network strength and residual factor scores by regressing the effects of age, sex, mean FD, scan sites, and mode-centered PDS for age 14. We used partial correlation to assess the association between residual network strength and residual TLFB and used an FDR correction (q < 0.05) for the multiple correlations.

Furthermore, we employed a three-wave bivariate latent change score model using the *lavvan* package in R and R studio to detect the linear change over time. This model allows us to quantify the longitudinal bidirectional influence between substance use and ICV over time (Nweze et al., 2023). Specifically, it facilitated an understanding of whether substance use predicted ICV and its brain activity, and vice versa. The key feature of this model is its ability to assess linear increases or decreases within the same construct across two adjacent waves. Change scores were calculated by regressing the observable score at a given timepoint from the previous timepoint (e.g., Δ Cig+CB in T1–T2 or Δ Cig+CB in T2–T3, where T1 = Timepoint 1, T2 = Timepoint 2, and T3 = Timepoint 3). Additionally, cross-lagged dynamic coupling (i.e., bidirectionality) was employed to explore individual differences in the relationships between substance use and linear changes in ICV/brain activity, as well as the relationship between ICV/brain activity and linear change in substance use. The model accounted for covariates such as age, sex and scan sites. For more details about the latent change score model, refer to the reference (Nweze et al., 2023).

As Fig. 6 shows, the latent change score model was specifically applied to examine the association between substance use and behaviours and brain activity associated with sustained attention. We focused on the relationship between the network strength of positive and negative networks, derived from Go and successful stop trials, and two types of substance use (Cig+CB and alcohol use). Notably, drug use data were excluded as adolescents at age 14 have no drug score. A total of 10 models were performed, and all model fit indices met the predefined criteria: CFI > 0.92, RMSEA <0.05, and SRMR < 0.03. An FDR correction (q < 0.05) was applied for multiple correlations. It is worth noting that all the correlations between substance use and sustained attention were conducted using the same sample across three timepoints.

#### 2.7.4 Permutation test

For the CPM analysis, we used a permutation test to assess the significance of the predictive performance, which is the correlation between the observed and predicted values. To generate a null distribution of these correlation values, we randomly shuffled the correspondence of the behavioural data and the PPI matrix of all subjects and reran the CPM pipeline with the shuffled data 1000 times. Based on this distribution, we set a threshold of *P <* 0.05 to determine the significance level at 95% for the predictive performance using 10-fold, 5-fold, and leave-site-out CV.

To estimate the significance of the predictive performance across timepoints and the external validation in the STRATIFY dataset, we shuffled the predictive values 1000 times. Then, we correlated the shuffled values with observed values to yield a null distribution of predictive correlation values. We also set a threshold of *P <* 0.05 to determine the significance level at 95% for the predictive performance across timepoints and generalization in STRATIFY.

## Acknowledgments

This work received support from the following sources: the European Union-funded FP6 Integrated Project IMAGEN (Reinforcement-related behaviour in normal brain function and psychopathology) (LSHM-CT-2007-037286), the Horizon 2020 funded ERC Advanced Grant ‘STRATIFY’ (Brain network based stratification of reinforcement-related disorders) (695313), Human Brain Project (HBP SGA 2, 785907, and HBP SGA 3, 945539), the Medical Research Council Grant ‘c-VEDA’ (Consortium on Vulnerability to Externalizing Disorders and Addictions) (MR/N000390/1), the National Institute of Health (NIH) (R01DA049238, A decentralized macro and micro gene-by-environment interaction analysis of substance use behaviour and its brain biomarkers), the National Institute for Health Research (NIHR) Biomedical Research Centre at South London and Maudsley NHS Foundation Trust and King’s College London, the Bundesministeriumfür Bildung und Forschung (BMBF grants 01GS08152; 01EV0711; Forschungsnetz AERIAL 01EE1406A, 01EE1406B; Forschungsnetz IMAC-Mind 01GL1745B), the Deutsche Forschungsgemeinschaft (DFG grants SM 80/7-2, SFB 940, TRR 265, NE 1383/14-1), the Medical Research Foundation and Medical Research Council (grants MR/R00465X/1 and MR/S020306/1), the National Institutes of Health (NIH) funded ENIGMA (grants 5U54EB020403-05 and 1R56AG058854-01), NSFC grant 82150710554 and European Union funded project ‘environMENTAL’, grant no: 101057429. Further support was provided by grants from: -the ANR (ANR-12-SAMA-0004, AAPG2019 -GeBra), the Eranet Neuron (AF12-NEUR0008-01 -WM2NA; and ANR-18-NEUR00002-01 -ADORe), the Fondation de France (00081242), the Fondation pour la Recherche Médicale (DPA20140629802), the Mission Interministérielle de Lutte-contre-les-Drogues-et-les-Conduites-Addictives (MILDECA), the Assistance-Publique-Hôpitaux-de-Paris and INSERM (interface grant), Paris Sud University IDEX 2012, the Fondation de l’Avenir (grant AP-RM-17-013), the Fédération pour la Recherche sur le Cerveau; the National Institutes of Health, Science Foundation Ireland (16/ERCD/3797), U.S.A. (Axon, Testosterone and Mental Health during Adolescence; RO1 MH085772-01A1) and by NIH Consortium grant U54 EB020403, supported by a cross-NIH alliance that funds Big Data to Knowledge Centres of Excellence.

ImagenPathways “Understanding the Interplay between Cultural, Biological and Subjective Factors in Drug Use Pathways” is a collaborative project supported by the European Research Area Network on Illicit Drugs (ERANID). This paper is based on independent research commissioned and funded in England by the National Institute for Health Research (NIHR) Policy Research Programme (project ref. PR-ST-0416-10001), China Scholarship Council—Trinity College Dublin Joint Scholarship Programme (202006750028). The views expressed in this article are those of the authors and not necessarily those of the national funding agencies or ERANID.

## Author contributions

R.W., J.K., L.R., and K.R. conceptualized this study. T.B., G.J.B., A.L.W.B., C.B., F.M.C., P.J.C., H.F., M.F.-B., J.G., H.G., P.G., A.H., B.I., K.M., J.-L.M., F.N., T.P., M.R., C.L., Z.P., M.-L.P.-M., M.N.S., A.S., M.R. and T.W.R. acquired the data. Y.W. processed and analysed the data. R.B., L.F., J.K., E.S., T.N. and J.H. contributed to data analyses and interpreted the results with Y.W. and R.W.. Y.W. and R.W. wrote the manuscript. All authors edited the paper and gave final approval before submission.

## Data and materials availability

IMAGEN data are available from a dedicated database: https://imagen2.cea.fr. Custom code that supports the findings of this study is available from the corresponding author upon request. All data needed to evaluate the conclusions in the paper are present in the paper and/or the Supplementary Materials. Additional data related to this paper may be requested from the authors.

