## Supplemental materials for "A robust brain network for sustained attention from adolescence to adulthood that predicts later substance use"

Yihe Weng *et al.*

**Supplementary Text**

***Method***

***Cambridge Neuropsychological Test Automated Battery (CANTAB)***

Several CANTAB tasks were used to examine cognitive abilities: the Affective Go/No-go task (AGN); the Rapid Visual Information Processing task (RVP), strategy working memory task (SWM), Cambridge guessing task (CGT) and strategy working memory (SWM). Only the subject at age 23 completed the CGT task. The RVP measures sustained attention. The AGN measures inhibitory control in the context of emotionally salient information, the CGT assesses impulsivity, the SWM assesses working memory. Detailed information on CANTAB was described in (Kuhn et al., 2020).

To examine the sustained attention network specificity, we correlated the network strength of predictive networks predicting ICV with CANTAB task at each timepoint separately. To control for potential confounders, we calculated residual network strength and residual performances of three CANTAB tasks by regressing the effects of age, sex, mean FD, scan sites, and mode-centered PDS for age 14. Finally, we used Spearman correlation to assess the association between residual sustained attention network strength and CANTAB performances.

To examine the specificity of sustained attention at baseline in influencing substance use, a two-sample t-test was performed to detect the significant difference in cigarette and cannabis use between high and lower cognition groups at baseline. Participants were categorized into 5 groups based on the ICV, network strength of positive and negative networks at age 14. The top of participants with the highest ICV/network strength of positive network, or lowest network strength of negative strength comprised the low sustained attention group, while the bottom of participants with lowest ICV/network strength of positive network, or highest network strength of negative strength constituted the high sustained attention group. Cig+CB were then compared between the higher and lower sustained attention groups at each time point. We found the significant coupling effect between Cig+CB and network strength derived from Go trials, instead of Successful stop trials. Here we only tested the difference in Cig+CB using positive and negative network derived from Go trials. We performed the similar analysis by stratifying the participants into higher SWM_BE group and lower SWM_BE group according to the between error value from the SWM task.

***Generalization in sub-groups from STRATIFY***

We tested if the predictive networks defined at age 23 in IMAGEN would generalize to distrinct patient groups in STRATIFY. STRATIFY includes several subgroups of individuals aged 23 with alcohol use disorder (AUD), major depression disorder (MDD), bulimia nervosa (BN), anorexia nervosa (AN) and 19 healthy controls.

***Dice coefficient***

We calculated the Dice coefficient (DC) to quantify the similarity of predictive networks across the three timepoints. A permutation test was also performed to estimate the predictive network similarity's significance. First, we shuffled the ICV at each timepoint and performed feature selection based on random behavioral phenotypes to yield random predictive networks, including positive and negative. Then we calculated DC from predictive networks between each pair of timepoint. These steps were iterated 1000 times to generate a null distribution of DC values. Finally, we set a threshold of *P* < 0.05 to determine the significance level at 95% for the similarity of the predictive networks between each timepoint.

***Comparison of predictive networks identified at one timepoint versus another***

Steiger’s Z value was employed to compare predictive performances of networks identified at different timepoints. This analysis involved comparing the R values derived from networks defined at distinct ages to predict ICV at the same age. For example, we compared the r values of brain networks defined at age 14 when predicting ICV at 19 (i.e., positive network: r = 0.25, negative network: r = 0.25, combined network: r = 0.28) with those R values of brain networks defined at age 19 itself (i.e., positive network: r = 0.16, negative network: r = 0.14, combined network: r = 0.16) derived from Go trials using Steiger's Z test (age 14 → age 19 vs. age 19 → 19). Similarly, comparisons were made between networks defined at age 14 predicting ICV at age 23 and those at age 23 predicting ICV at age 23 (age 14 → age 23 vs. age 23 → 23), as well as between networks defined at age 19 predicting ICV at age 23 and those at age 23 predicting ICV at age 23 (age 19 → age 23 vs. age 23 → age 23). These comparisons were performed separately for Go trials and Successful Stop trials.

***CPM analysis using Failed stop trials***

We performed another CPM analysis using Failed stop trials using gPPI matrix obtained from the second GLM, described in the main text. The CPM analysis was conducted using 10-fold CV, 5-fold CV and leave-site-out CV.

***Prediction across timepoints controlling for ICV at age 14***

To examine whether connectivity predictors shared variations of sustained attention across timepoints, we applied predictive models developed at ages 14 and 19 to predict ICV at subsequent timepoints controlling for ICV at age 14. Specifically, we used predictive models (including parameters and selected edges) developed at age 14 to predict ICV at ages 19 and 23 separately. First, we calculated the network strength using the gPPI matrix at ages 19 and 23 based on the selected edges identified from CPM analysis at age 14. We then estimated the predicted ICV at ages 19 and 23 by applying the linear model parameters (slope and intercept) obtained from CPM analysis at age 14 to the network strength. Finally, we evaluated the predictive performance by calculating the partial correlation between the predicted and observed values at ages 19 and 23, controlling for ICV at age 14. Similarly, we applied models developed at age 19 to predict ICV at age 23, also controlling for ICV at age 14. To assess the significance of the predictive performance, we used a permutation test, shuffling the predicted ICV values and calculating partial correlation to general a random distribution over 1,000 iterations.

***Results***

***Specificity of sustained attention network***

Sustained attention network strength derived from Go trials and Successful stop trials was significantly correlated with RVPA (all *P* < 0.05) for both negative and positive networks at ages 14 and 19 but not with AGN task performance (all *P* > 0.05) (Table S12). These results suggest that the networks derived from Go trails and Successful stop trials are specific to sustained attention.

No significant difference in Cig+CB was found between high and low sustained attention groups (obtained from both behavior level and brain activity) at age 14 (all *P* > 0.462). Higher Cig+CB use was found in low sustained attention group compared to high sustained attention group at age 19 (all *P* < 0.021) and age 23 (all *P* < 0.007) (Table S22). In addition, no significant difference in Cig+CB was found at ages 14 (t = 0.11, *P* = 0.912), 19 (t = 1.65, *P* = 0.10), and 23 (t = 1.43, *P* = 0.154) between higher and lower SWM_BE groups (Table S23).

***CPM predictive performance under Failed stop trials***

Positive, negative, and combined networks derived from Failed stop trials significantly predicted ICV: at age 14 (r = 0.10, *P* = 0.033; r = 0.19, *P* < 0.001; and r = 0.17, *P* < 0.001, respectively), at age 19 (r = 0.21; r = 0.18; and r = 0.21, all *P* < 0.001, respectively), and at age 23 (r = 0.33, r = 0.35, and r = 0.36, respectively, all *P* < 0.001). We obtained similar results using a 5-fold CV and leave-site-out CV (Table S6).

***Predictive network similarity***

With respect to Go trials, the mean DC values for the positive and negative networks across all three timepoints were 0.06 and 0.03, respectively, for ICV. With respect to Successful stop trials, the mean DC values for positive and negative networks predicting ICV across all three timepoints were 0.01 and 0.01.

Positive and negative networks predicting ICV derived from Go trials were significantly similar between ages 14 and 19 (DC = 0.03, *P* = 0.001 and DC = 0.03, *P* < 0.001), and between ages 19 and 23 (DC = 0.06 and DC = 0.04, respectively, all *P* < 0.001) (FDR correction, 0.05). The positive network predicting ICV derived from Go trials was significantly similar between ages 14 and 23 (DC = 0.07, *P* < 0.001) (FDR correction, 0.05). The mean DC of the positive and negative networks predicting ICV across 3 timepoints derived from Go trials are 0.06 and 0.03 respectively.

The negative networks predicting ICV derived from Successful stop trials were significantly similar between ages 14 and 19 (DC = 0.03, *P* = 0.001) (FDR correction, q < 0.05). The mean DC of the positive and negative networks predicting ICV derived from Successful stop trials was 0.01 and 0.01. Detailed results about DC between each pair of timepoints can be found in Table S13.

***Generalization in sub-groups in STRATIFY***

We examined generalization to separate patient cohorts in STRATIFY. Brain networks predicting ICV derived from Go trials defined at age 23 generalized to almost all patient cohorts, including AUD, MDD, BN, and AN (all *P* < 0.05). The prediction for the healthy controls was moderately accurate (r ~ 0.4), although this was not statistically significant due to the small sample size (n=19) for Go trials (Fig. S8A, left panel). However, brain networks predicting ICV derived from Successful stop trials failed to predict ICV in individuals with AUD (*P* > 0.05), although they generalized to other patient groups (Fig. S8A, right panel). Furthermore, the correlations between sustained attention network strength of positive, negative, and combined networks derived from Successful stop trials and ICV in the groups with AUD were in the opposite direction compared with all other groups (Fig. S8B).

***Comparison of predictive performance at different timepoints***

Steiger’s Z value was used to test if the difference in R values obtained using predictive networks at one timepoint versus another. For positive, negative, and combined networks predicting ICV derived from Go trials at age 19, the R values were higher when using predictive networks defined at 19 than those defined at 14 (Z = 3.79, Z = 3.39, Z = 3.99, all *P* < 0.00071). Similarly, the R values for positive, negative, and combined networks predicting ICV derived from Go trials at age 23 were higher when using predictive networks defined at age 23 compared to those defined at ages 14 (Z = 6.00, Z = 5.96, Z = 6.67, all *P* < 3.47e^-9^) or 19 (Z = 2.80, Z = 2.36, Z = 2.57, all *P* < 0.005).

At age 19, the R value for the positive network predicting ICV derived from Successful stop trials was higher when using predictive networks defined at 19 compared to those defined at 14 (Z = 1.54, *P* = 0.022), while the negative and combined networks did not show a significant difference (Z = 0.85, P = 0.398; Z = 2.29, *P* = 0.123). At age 23, R values for the positive and combined networks predicting ICV derived from Successful stop trials were higher when using predictive networks defined at 23 compared to those defined at 14 (Z = 3.00, Z = 2.48, all *P* < 3.47e^-9^) or 19 (Z = 2.52, Z = 1.99, all *P* < 0.005). However, the R value for the negative network at age 23 did not significantly differ when using predictive networks defined at 14 (Z = 1.80, *P* = 0.072) or 19 (Z = 1.48, *P* = 0.138).

***Correlation between drug use and behavior and brain activity***

ICV negatively correlated with drug use at age 19 (rho = -0.11, *P* =0.001) (Table S14). ICV at age 23 negatively correlated with drug use at age 19 (rho = -0.08, P =0.014) (Table S17). Sustained attention network strength derived from Successful stop trials significantly correlated with drug use at age 19 for the positive network (rho = -0.12, *P* < 0.001; FDR correction, 0.05) (Table S18). Sustained attention network strength derived from Successful stop trials at age 23 correlated with drug use at age 19 (Positive network: rho = -0.09, *P* =0.005; Negative network: rho = 0.09, *P* =0.007) (Table S21).

***Predictions across timepoints controlling for ICV at age 14***

Positive and combined networks derived from Go trials defined at age 14 predicted ICV at ages 19 (r = 0.10, *P* = 0.028; r = 0.08, *P* = 0.047) but negative network did not (r = 0.06, *P* = 0.119). Positive network derived from Go trials defined at age 14 predicted ICV at age 23 (r = 0.11, *P* = 0.013) but negative and combined networks did not (r = 0.04, *P* = 0.187; r = 0.08, *P* = 0.056). Positive, negative, and combined networks derived from Go trials defined at age 19 predicted ICV at age 23 (r = 0.22, r = 0.19, and r = 0.22, respectively, all *P* < 0.001).

Positive, negative, and combined networks derived from Successful stop trials defined at age 14 predicted ICV at age 19 (r = 0.08, *P* = 0.036; r = 0.10, *P* = 0.012; r = 0.11, *P* = 0.009) and 23 (r = 0.11, *P* = 0.005; r = 0.13, *P* = 0.005; r = 0.13, *P* = 0.017) respectively. Positive, negative, and combined networks derived from Successful stop trials defined at age 19 predicted ICV at age 23 (r = 0.18, r = 0.18, and r = 0.17, respectively, all *P* < 0.001).

***Discussion***

***Sustained attention networks functioning as global activation***

Our findings strongly indicate that sustained attention relies on global brain activation (i.e., network strength) rather than specific regions or networks (see also (Zhao et al., 2021)). We observed brain networks associated with high- or low-sustained attention span in large-scale networks across the cortex, subcortex, and cerebellum across adolescence to adulthood (See Figs. S2-3, consistent with (Rosenberg et al., 2020)), instead of being confined to a few key regions. In our study, however, although the edges in the sustained attention networks were significantly similar from ages 14 to 23 (Table S13), there were relatively few overlapping edges in the predictive networks over time. It is worth noting that Dice coefficient values depend heavily on the significance threshold applied to the data (Frohner, Teckentrup, Smolka, & Kroemer, 2019). However, sustained attention network patterns identified could efficiently predict sustained attention for the subsequent timepoints. A prior study (Cai et al., 2019) has shown that children aged 9-12 could recruit key nodes (e.g., rIFG and rMFG), eliciting an adult-like global activation pattern that predicted their inhibitory control abilities. Similarly, our findings suggest that adolescents likely exhibit adult-like global activation patterns predicting sustained attention.

***Aberrant sustained attention network in AUD***

A notable exception was the failure of the network derived from Successful stop trials to generalize to patients with AUD, requiring higher attention levels. We speculate that activity in the sustained attention network of individuals with AUD might be similar to healthy adults during low cognitive demands but abnormal compared to healthy adults when faced with higher cognitive demands. Evidence from past literature shows that alcohol misuse is associated with attention deficits and dysfunctional neural mechanisms (Gunn, Mackus, Griffin, Munafo, & Adams, 2018; Li, Chen, Tang, & Li, 2021; Narayan, Aitken, Downey, & Hayley, 2021; Spear, 2018). Furthermore, lower activation of parietal and prefrontal cortices has been observed in abstinent patients with AUD compared with healthy controls during visual attention tasks (Zehra et al., 2019), suggesting that differences in the attention network of individuals with AUD might underlie attention deficits in AUD.

Although the sustained attention network derived from Successful stop trials seen in healthy controls failed to generalize to AUD, we observed no significant difference in behavioral measures – ICV – between healthy controls and those with AUD. Our results may reflect compensatory mechanisms in AUD that allow these individuals to complete sustained attention tasks, which is consistent with prior studies (Tapert et al., 2004; Zehra et al., 2019). Compensation manifests as abnormal brain activity while performing normally on the task (Chanraud, Pitel, Muller-Oehring, Pfefferbaum, & Sullivan, 2013). Zehra et al. (Zehra et al., 2019) found brain activation differences during attention tasks despite no differences in behavioral performance between the abstinent patients with AUD and healthy controls. Previous studies (Chanraud et al., 2013; Squeglia, Jacobus, & Tapert, 2009) pointed out that individuals with AUD might exhibit subtle neural reorganization or compensation to preserve normal cognitive abilities. Tapert et al. (Tapert et al., 2004) found that heavy and light drinkers had similar behavioral performance on a working memory task. Activation differences were found in the parietal and occipital lobes and the cerebellar, indicating subtle neuronal reorganization may occur in AUD. Similarly, a study found that individuals with AUD maintained standard working memory by recruiting other cerebellar-based functional networks to complete the task (Chanraud et al., 2013). Successful completion of working memory tasks requires sustained attention (Myers, Stokes, & Nobre, 2017). These studies suggest that the failure of the sustained attention network derived from Successful stop trials to generalize AUD in the current study may be due to compensatory mechanisms employed by individuals with AUD while completing tasks requiring high-level sustained attention.

***Specificity of the prediction of predictive networks***

We found that task-related function connectivity derived from Go trials, Successful stop trials, and Failed stop trials successfully predicted sustained attention across three timepoints. However, predictive performances of predictive networks derived from Go trials were higher than those derived from Successful stop trials and Failed stop trials. These results suggest that sustained attention is particularly crucial during Go trials when participants need to respond to the Go signal. In contrast, although Successful Stop and Failed Stop trials also require sustained attention, these tasks primarily involve inhibitory control along with sustained attention.

***Reference***

Cai, W., Duberg, K., Padmanabhan, A., Rehert, R., Bradley, T., Carrion, V., & Menon, V. (2019). Hyperdirect insula-basal-ganglia pathway and adult-like maturity of global brain responses predict inhibitory control in children. *Nature Communications, 10*(1), 4798. doi:10.1038/s41467-019-12756-8

Chanraud, S., Pitel, A. L., Muller-Oehring, E. M., Pfefferbaum, A., & Sullivan, E. V. (2013). Remapping the brain to compensate for impairment in recovering alcoholics. *Cereb Cortex, 23*(1), 97-104. doi:10.1093/cercor/bhr381

Frohner, J. H., Teckentrup, V., Smolka, M. N., & Kroemer, N. B. (2019). Addressing the reliability fallacy in fMRI: Similar group effects may arise from unreliable individual effects. *Neuroimage, 195*, 174-189. doi:10.1016/j.neuroimage.2019.03.053

Gunn, C., Mackus, M., Griffin, C., Munafo, M. R., & Adams, S. (2018). A systematic review of the next-day effects of heavy alcohol consumption on cognitive performance. *Addiction, 113*(12), 2182-2193. doi:10.1111/add.14404

Kuhn, S., Lisofsky, N., Banaschewski, T., Barker, G., Bokde, A. L. W., Bromberg, U., . . . Consortium, I. (2020). Hierarchical associations of alcohol use disorder symptoms in late adolescence with markers during early adolescence. *Addict Behav, 100*, 106130. doi:10.1016/j.addbeh.2019.106130

Li, G., Chen, Y., Tang, X., & Li, C. R. (2021). Alcohol use severity and the neural correlates of the effects of sleep disturbance on sustained visual attention. *J Psychiatr Res, 142*, 302-311. doi:10.1016/j.jpsychires.2021.08.018

Myers, N. E., Stokes, M. G., & Nobre, A. C. (2017). Prioritizing Information during Working Memory: Beyond Sustained Internal Attention. *Trends in Cognitive Sciences, 21*(6), 449-461. doi:10.1016/j.tics.2017.03.010

Narayan, A. J., Aitken, B., Downey, L. A., & Hayley, A. C. (2021). The effects of amphetamines alone and in combination with alcohol on functional neurocognition: A systematic review. *Neurosci Biobehav Rev, 131*, 865-881. doi:10.1016/j.neubiorev.2021.10.003

Rosenberg, M. D., Scheinost, D., Greene, A. S., Avery, E. W., Kwon, Y. H., Finn, E. S., . . . Chun, M. M. (2020). Functional connectivity predicts changes in attention observed across minutes, days, and months. *Proc Natl Acad Sci U S A, 117*(7), 3797-3807. doi:10.1073/pnas.1912226117

Spear, L. P. (2018). Effects of adolescent alcohol consumption on the brain and behaviour. *Nature Reviews Neuroscience, 19*(4), 197-214. doi:10.1038/nrn.2018.10

Squeglia, L. M., Jacobus, J., & Tapert, S. F. (2009). The influence of substance use on adolescent brain development. *Clin EEG Neurosci, 40*(1), 31-38. doi:10.1177/155005940904000110

Tapert, S. F., Schweinsburg, A. D., Barlett, V. C., Brown, S. A., Frank, L. R., Brown, G. G., & Meloy, M. J. (2004). Blood oxygen level dependent response and spatial working memory in adolescents with alcohol use disorders. *Alcohol Clin Exp Res, 28*(10), 1577-1586. doi:10.1097/01.alc.0000141812.81234.a6

Zehra, A., Lindgren, E., Wiers, C. E., Freeman, C., Miller, G., Ramirez, V., . . . Volkow, N. D. (2019). Neural correlates of visual attention in alcohol use disorder. *Drug Alcohol Depend, 194*, 430-437. doi:10.1016/j.drugalcdep.2018.10.032

Zhao, W., Palmer, C. E., Thompson, W. K., Chaarani, B., Garavan, H. P., Casey, B. J., . . . Fan, C. C. (2021). Individual Differences in Cognitive Performance Are Better Predicted by Global Rather Than Localized BOLD Activity Patterns Across the Cortex. *Cereb Cortex, 31*(3), 1478-1488. doi:10.1093/cercor/bhaa290

**Table S1 Demographic information for CPM analysis.**

|  | **Age 14** | **Age 19** | **Age 23** | **STRATIFY** |
| --- | --- | --- | --- | --- |
| N | 716 | 1079 | 1120 | 304 |
| Age (Years) | 14.6 ± 0.4 | 19.1 ± 0.8 | 22.6 ± 0.7 | 22 ± 2.2 |
| Sex (M/F) | 316/400 | 496/583 | 530/590 | 240/64 |
| Site | 119/103/47/51/  121/82/90/103 | 145/152/76/98/  159/128/173/148 | 144/163/123/149/  151/143/123/124 | 24/0/129/151 |
| Mean FD | 0.19 ± 0.1 | 0.15 ± 0.08 | 0.17 ± 0.08 | 0.17 ± 0.09 |
| ICV | 0.234±0.038 | 0.222±0.049 | 0.217±0.054 | 0.21 ± 0.05 |
| GO RT | 430.4 ± 61.6 | 397 ± 68.4 | 402.6 ± 73.2 | 427.8 ± 72.4 |
| SSD | 225.3 ± 84.4 | 173.5 ± 91.3 | 181.3 ± 122.5 | 180.2 ± 97.9 |
| Stop RT | 404 ± 69.8 | 353.9 ± 72.4 | 362.9 ± 78.7 | 376.1 ± 66.4 |
| pOmission | 1.3% ± 2.6% | 1.8% ± 4.6% | 3.1% ± 8.7% | 3% ± 8.6% |
| pChoiceError | 5.4% ± 7.7% | 4.7% ± 3.8% | 5.2% ± 7.7% | 5.3% ± 4.7% |
| pCommission | 49.6% ± 1.6% | 48.3% ± 3.8% | 47.5% ± 6% | 48% ± 5.1% |

**Note:** N, number of subjects; Site, London/ Nottingham/ Dublin/ Berlin/ Hamburg/ Mannheim/ Paris/ Dresden; ICV, intra-individual coefficient of variation; GO RT, reaction time in Go trials; SSD, stop signa delay; Stop RT, reaction time in Stop trials; pOmission, probability of go omissions (no response); pChoiceError, probability of choice errors derived from Go trialss; pCommission, probability of commission on Stop trials.

**Table S2 Demographic information across three timepoints for exploratory factor analysis.**

|  | Age 14 | Age 19 | Age 23 |
| --- | --- | --- | --- |
| N | 2047 | 1386 | 1166 |
| Age (Years) | 14.4 ± 0.4 | 19 ± 0.7 | 22.6 ± 0.7 |
| Sex (M/F) | 989/1027 | 664/721 | 550/614 |
| Site | 254/328/209/256/  262/246/248/213 | 191/224/147/97/  155/178/209/184 | 162/185/138/156/  167/147/60/149 |
| Mean FD | 0.28 ± 0.29 | 0.18 ± 0.13 | 0.18 ± 0.12 |
| GO RT | 467.5 ± 80 | 399.6 ± 70.4 | 405.4 ± 73.2 |
| SSD | 320.4 ± 148.3 | 187 ± 130.6 | 191.2 ± 162.5 |
| Stop RT | 462.4 ± 114.4 | 359 ± 80.2 | 364.8 ± 77.7 |
| pOmission | 4.5% ± 10.7% | 2.6% ± 8.5% | 3.9% ± 11.4% |
| pChoiceError | 4.7% ± 6.8% | 4.7% ± 4.7% | 5.3% ± 7.8% |
| pCommission | 47.9% ± 6.4% | 47.5% ± 6% | 47.1% ± 7.1% |

**Note:** N, number of subjects; Site, London/ Nottingham/ Dublin/ Berlin/ Hamburg/ Mannheim/ Paris/ Dresden; GO RT, reaction time in Go trials; SSD, stop signa delay; Stop RT, reaction time in Stop trials; pOmission, probability of go omissions (no response); pChoiceError, probability of choice errors derived from Go trialss; pCommission, probability of commission on Stop trials.

**Table S3 Linear mixed model of intra-individual coefficient of variation (ICV).**

|  | Sum (Sq) | Mean (Sq) | NumDF | DenDF | F value | *Pr*(>F) |
| --- | --- | --- | --- | --- | --- | --- |
| Timepoint | 59.8 | 29.9 | 2 | 1895.3 | 51.1449 | <0.001 |
| Sex | 1.022 | 1.0223 | 1 | 1483.1 | 1.7486 | 0.186 |
| Site | 28.527 | 4.0753 | 7 | 1489.4 | 6.971 | <0.001 |
| Mode-center PDS | 0.887 | 0.8875 | 1 | 1488.8 | 1.518 | 0.218 |
| Age | 0.002 | 0.0019 | 1 | 1464.1 | 0.0032 | 0.955 |

**Table S4 Post-hoc test for intra-individual coefficient of variation (ICV).**

| Contrast | Estimate | SE | DF | t | *P* value |
| --- | --- | --- | --- | --- | --- |
| Age 14 - Age 19 | 0.263 | 0.040 | 2093 | 6.535 | <.0001 |
| Age 14 - Age 23 | 0.423 | 0.042 | 2060 | 10.109 | <.0001 |
| Age 19 - Age 23 | 0.159 | 0.033 | 1688 | 4.768 | <.0001 |

**Table S5 Correlation of intra-individual coefficient of variation (ICV) between timepoints.**

| Correlation | N | R value | *P* value |
| --- | --- | --- | --- |
| Age 14 - Age 19 | 453 | 0.35 | <2.5e^-14^ |
| Age 14 - Age 23 | 418 | 0.32 | <3.2e^-11^ |
| Age 19 - Age 23 | 756 | 0.45 | <1e^-16^ |

**Table S6 Predictive performance for intra-individual coefficient of variation (ICV) from various cross-validation (CV) schemes across three timepoints derived from Go trials, Successful stop trials and Failed stop trials.**

| Condition | Timepoint | Predictive network | 10-fold CV | | | 5-fold CV | | | leave-site-out CV | | |
| --- | --- | --- | --- | --- | --- | --- | --- | --- | --- | --- | --- |
|  |  |  | r | *P* | R² (%) | r | *P* | R² (%) | r | *P* | R² (%) |
| Go trials | Age 14 | Positive | 0.25 | <0.001 | 6.34 | 0.24 | <0.001 | 5.81 | 0.26 | <0.001 | 6.71 |
|  |  | Negative | 0.25 | <0.001 | 6.46 | 0.24 | <0.001 | 5.89 | 0.26 | <0.001 | 6.75 |
|  |  | Combined | 0.28 | <0.001 | 8.07 | 0.27 | <0.001 | 7.43 | 0.30 | <0.001 | 8.82 |
|  | Age 19 | Positive | 0.27 | <0.001 | 7.42 | 0.27 | <0.001 | 7.11 | 0.19 | <0.001 | 3.43 |
|  |  | Negative | 0.25 | <0.001 | 6.42 | 0.25 | <0.001 | 6.09 | 0.17 | <0.001 | 2.86 |
|  |  | Combined | 0.28 | <0.001 | 7.90 | 0.28 | <0.001 | 7.60 | 0.20 | <0.001 | 3.81 |
|  | Age 23 | Positive | 0.38 | <0.001 | 14.26 | 0.38 | <0.001 | 14.41 | 0.26 | <0.001 | 6.76 |
|  |  | Negative | 0.33 | <0.001 | 11.00 | 0.33 | <0.001 | 10.87 | 0.18 | <0.001 | 3.20 |
|  |  | Combined | 0.37 | <0.001 | 14.07 | 0.38 | <0.001 | 14.17 | 0.23 | <0.001 | 5.34 |
| Successful stop | Age 14 | Positive | 0.22 | <0.001 | 4.82 | 0.19 | <0.001 | 3.69 | 0.15 | <0.001 | 2.25 |
|  |  | Negative | 0.12 | 0.017 | 1.39 | 0.11 | 0.012 | 1.23 | 0.07 | 0.028 | 0.46 |
|  |  | Combined | 0.20 | <0.001 | 4.16 | 0.18 | <0.001 | 3.27 | 0.13 | <0.001 | 1.65 |
|  | Age 19 | Positive | 0.19 | <0.001 | 3.79 | 0.20 | <0.001 | 4.04 | 0.11 | <0.001 | 4.22 |
|  |  | Negative | 0.15 | 0.001 | 2.21 | 0.15 | <0.001 | 2.32 | 0.09 | 0.002 | 4.26 |
|  |  | Combined | 0.18 | <0.001 | 3.31 | 0.19 | <0.001 | 3.60 | 0.11 | <0.001 | 1.14 |
|  | Age 23 | Positive | 0.24 | <0.001 | 5.86 | 0.24 | <0.001 | 6.02 | 0.14 | <0.001 | 1.95 |
|  |  | Negative | 0.21 | <0.001 | 4.43 | 0.21 | <0.001 | 4.54 | 0.14 | <0.001 | 2.06 |
|  |  | Combined | 0.23 | <0.001 | 5.33 | 0.24 | <0.001 | 5.58 | 0.15 | <0.001 | 2.22 |
| Failed stop | Age 14 | Positive | 0.10 | 0.033 | 0.98 | 0.11 | 0.009 | 1.20 | 0.11 | 0.001 | 1.17 |
|  |  | Negative | 0.19 | <0.001 | 3.55 | 0.18 | <0.001 | 3.16 | 0.16 | <0.001 | 2.50 |
|  |  | Combined | 0.17 | <0.001 | 3.04 | 0.17 | <0.001 | 3.04 | 0.16 | <0.001 | 2.68 |
|  | Age 19 | Positive | 0.21 | <0.001 | 4.42 | 0.20 | <0.001 | 4.10 | 0.16 | <0.001 | 2.69 |
|  |  | Negative | 0.18 | <0.001 | 3.33 | 0.18 | <0.001 | 3.35 | 0.13 | <0.001 | 1.82 |
|  |  | Combined | 0.21 | <0.001 | 4.45 | 0.21 | <0.001 | 4.34 | 0.17 | <0.001 | 2.79 |
|  | Age 23 | Positive | 0.33 | <0.001 | 11.09 | 0.33 | <0.001 | 10.55 | 0.21 | <0.001 | 4.57 |
|  |  | Negative | 0.35 | <0.001 | 12.20 | 0.34 | <0.001 | 11.63 | 0.22 | <0.001 | 4.98 |
|  |  | Combined | 0.36 | <0.001 | 12.64 | 0.35 | <0.001 | 12.04 | 0.23 | <0.001 | 5.22 |

**Table S7 Correlation of network strength of sustained attention network between timepoints.**

| Correlation | Predictive networks | Go trials |  | Successful stop trials | |
| --- | --- | --- | --- | --- | --- |
|  |  | R value | *P* value | R value | *P* value |
| Age 14 - Age 19 | positive | 0.20 | 1.32E-05 | 0.15 | 0.001 |
|  | negative | 0.23 | 1.25E-06 | 0.20 | 5.48E-05 |
|  | combined | 0.31 | <0.001 | 0.20 | 4.36E-08 |
| Age 14 - Age 23 | positive | 0.14 | 0.003 | 0.21 | 8.24E-06 |
|  | negative | 0.19 | 8.56E-05 | 0.18 | 2.14E-04 |
|  | combined | 0.21 | 3.72E-09 | 0.21 | 7.79E-09 |
| Age 19 - Age 23 | positive | 0.18 | 1.58E-04 | 0.21 | 4.93E-06 |
|  | negative | 0.25 | 1.35E-07 | 0.25 | 2.35E-07 |
|  | combined | 0.28 | 2.22E-15 | 0.23 | 1.53E-10 |

**Table S8 Predictive performance of intra-individual coefficient of variation (ICV) at each timepoint with distinct feature-selection thresholds using 10-fold cross validation.**

| Timepoint | P value | Go trials | | | Successful stop trials | | |
| --- | --- | --- | --- | --- | --- | --- | --- |
|  |  | Positive | Negative | Combined | Positive | Negative | Combined |
| Age 14 | 0.1 | 0.22 | 0.22 | 0.26 | 0.16 | 0.15 | 0.18 |
|  | 0.05 | 0.23 | 0.23 | 0.26 | 0.16 | 0.12 | 0.16 |
|  | 0.01 | 0.25 | 0.25 | 0.28 | 0.22 | 0.11 | 0.2 |
|  | 0.005 | 0.25 | 0.27 | 0.29 | 0.21 | 0.12 | 0.2 |
|  | 0.001 | 0.22 | 0.25 | 0.27 | 0.14 | 0.14 | 0.16 |
|  | 0.0005 | 0.21 | 0.22 | 0.25 | 0.12 | 0.14 | 0.14 |
|  | 0.0001 | 0.2 | 0.17 | 0.21 | 0.14 | 0.11 | 0.13 |
| Age 19 | 0.1 | 0.29 | 0.28 | 0.3 | 0.24 | 0.17 | 0.21 |
|  | 0.05 | 0.29 | 0.27 | 0.29 | 0.23 | 0.17 | 0.22 |
|  | 0.01 | 0.27 | 0.25 | 0.28 | 0.2 | 0.15 | 0.18 |
|  | 0.005 | 0.26 | 0.24 | 0.27 | 0.19 | 0.14 | 0.17 |
|  | 0.001 | 0.24 | 0.18 | 0.22 | 0.2 | 0.15 | 0.17 |
|  | 0.0005 | 0.24 | 0.17 | 0.2 | 0.19 | 0.17 | 0.17 |
|  | 0.0001 | 0.26 | 0.2 | 0.22 | 0.19 | 0.23 | 0.19 |
| Age 23 | 0.1 | 0.43 | 0.4 | 0.42 | 0.27 | 0.22 | 0.26 |
|  | 0.05 | 0.42 | 0.38 | 0.42 | 0.26 | 0.23 | 0.25 |
|  | 0.01 | 0.38 | 0.33 | 0.37 | 0.24 | 0.22 | 0.23 |
|  | 0.005 | 0.36 | 0.32 | 0.36 | 0.26 | 0.23 | 0.25 |
|  | 0.001 | 0.36 | 0.3 | 0.35 | 0.29 | 0.28 | 0.28 |
|  | 0.0005 | 0.37 | 0.29 | 0.36 | 0.28 | 0.3 | 0.28 |
|  | 0.0001 | 0.38 | 0.29 | 0.36 | 0.3 | 0.33 | 0.3 |

**Table S9 Prediction for intra-individual coefficient of variation (ICV) across timepoint.**

| Predictive models | Predictive networks | Go trials |  | Successful stop trials | |
| --- | --- | --- | --- | --- | --- |
|  |  | r value | *P* value | r value | *P* value |
| Age 14 predicts Age 19 | positive | 0.16 | <0.001 | 0.11 | <0.001 |
|  | negative | 0.14 | 0.001 | 0.12 | <0.001 |
|  | combined | 0.16 | <0.001 | 0.13 | <0.001 |
| Age 14 predicts Age 23 | positive | 0.20 | <0.001 | 0.14 | <0.001 |
|  | negative | 0.12 | <0.001 | 0.15 | <0.001 |
|  | combined | 0.17 | <0.001 | 0.15 | <0.001 |
| Age 19 predicts Age 23 | positive | 0.30 | <0.001 | 0.17 | <0.001 |
|  | negative | 0.26 | <0.001 | 0.16 | <0.001 |
|  | combined | 0.31 | <0.001 | 0.17 | <0.001 |
| Age 23 predicts STRATIFY | positive | 0.34 | <0.001 | 0.26 | <0.001 |
|  | negative | 0.34 | <0.001 | 0.22 | <0.001 |
|  | combined | 0.35 | <0.001 | 0.26 | <0.001 |

**Table S10 Predictive performance of ICV in patients’ groups in STRATIFY.**

| Groups | N | R _positive_ | *P* _positive_ | R _negative_ | *P* _negative_ | R _combined_ | *P* _combined_ |
| --- | --- | --- | --- | --- | --- | --- | --- |
| *Go trials* |  |  |  |  |  |  |  |
| Control | 19 | 0.38 | 0.107 | 0.40 | 0.088 | 0.39 | 0.096 |
| AUD | 79 | 0.22 | 0.038 | 0.36 | 0.001 | 0.29 | 0.007 |
| MDD | 87 | 0.36 | <0.001 | 0.25 | 0.012 | 0.33 | 0.001 |
| BN | 42 | 0.41 | 0.006 | 0.43 | 0.004 | 0.44 | 0.004 |
| AN | 52 | 0.49 | <0.001 | 0.53 | <0.001 | 0.52 | <0.001 |
| *Successful stop trials* | | | | | | | |
| Control | 19 | 0.46 | 0.045 | 0.27 | 0.269 | 0.46 | 0.046 |
| AUD | 79 | 0.07 | 0.516 | 0.10 | 0.364 | 0.06 | 0.571 |
| MDD | 87 | 0.23 | 0.033 | 0.30 | 0.005 | 0.28 | 0.009 |
| BN | 42 | 0.32 | 0.040 | 0.24 | 0.127 | 0.30 | 0.051 |
| AN | 52 | 0.42 | 0.002 | 0.27 | 0.056 | 0.37 | 0.008 |

Note: AUD, alcohol use disorder; MDD, major depression disorder; BN, bulimia Nervosa; AN, anorexia nervosa. N, number of subjects.

**Table S11 Timeline Followback items used for exploratory factor analysis.**

| ID | Item | Content |
| --- | --- | --- |
| 1 | tlfb_alcohol1 | Total of days using alcohol in the past 30 days |
| 2 | tlfb_alcohol2 | Total of alcohol drink units in past 30 days |
| 3 | tlfb_alcohol3 | Total of days using = 5 (for boys)/ = 4 (for girls) alcohol drink units in past 30 days |
| 4 | tlfb_alcohol4 | Total cost of alcohol in past 30 days: |
| 5 | tlfb_cannabis1 | Total of days using cannabis in the past 30 days |
| 6 | tlfb_cocaine1 | Total of days using cocaine in the past 30 days |
| 7 | tlfb_ecstasy1 | Total of days using ecstasy in the past 30 days |
| 8 | tlfb_ketamine1 | Total of days using ketamine in the past 30 days |
| 9 | tlfb_tobacco1 | Total of days smoking tobacco in the past 30 days |

Note: Factor analysis applied on age 14 used items 1-6 because no adolescents at age 14 used drugs, while factor analysis applied on ages 19 and 23 used items 1-9.

**Table S12 Correlation between sustained attention network strength and CANTAB test.**

| Correlation | Predictive network | Successful stop trials | | Go trials |  |
| --- | --- | --- | --- | --- | --- |
|  |  | r value | *P* value | r value | *P* value |
| age 14 strength & age 14 RVPA | positive | **-0.09** | **0.022** | **-0.10** | **0.006** |
|  | negative | **0.10** | **0.007** | **0.08** | **0.028** |
| age 19 strength & age 19 RVPA | positive | **-0.14** | **<0.001** | **-0.15** | **<0.001** |
|  | negative | **0.16** | **<0.001** | **0.13** | **<0.001** |
| age 23 strength & age 23 RVPA | positive | — | — | — | — |
|  | negative | — | — | — | — |
| age 14 strength & age 14 AGN_POS | positive | -0.03 | 0.440 | -0.07 | 0.069 |
|  | negative | 0.05 | 0.206 | 0.03 | 0.435 |
| age 19 strength & age 19 AGN_POS | positive | -0.04 | 0.188 | -0.06 | 0.070 |
|  | negative | 0.04 | 0.152 | 0.04 | 0.146 |
| age 23 strength & age 23 AGN_POS | positive | — | — | — | — |
|  | negative | — | — | — | — |
| age 14 strength & age 14 AGN_NEG | positive | -0.05 | 0.212 | -0.06 | 0.112 |
|  | negative | 0.06 | 0.148 | 0.01 | 0.747 |
| age 19 strength & age 19 AGN_NEG | positive | -0.01 | 0.744 | -0.02 | 0.563 |
|  | negative | 0.02 | 0.495 | 0.02 | 0.488 |
| age 23 strength & age 23 AGN_NEG | positive | — | — | — | — |
|  | negative | — | — | — | — |
| age 14 strength & age 14 CGT_DA | positive | 0.05 | 0.250 | 0.08 | 0.058 |
|  | negative | -0.03 | 0.430 | -0.06 | 0.137 |
| age 19 strength & age 19 CGT_DA | positive | 0.05 | 0.083 | **0.07** | **0.015** |
|  | negative | **-0.07** | **0.016** | **-0.12** | **<0.001** |
| age 23 strength & age 23 CGT_DA | positive | 0.05 | 0.076 | 0.05 | 0.115 |
|  | negative | **-0.07** | **0.014** | **-0.06** | **0.031** |
| age 14 strength & age 14 CGT_RT | positive | 0.05 | 0.227 | 0.00 | 0.910 |
|  | negative | 0.03 | 0.454 | 0.02 | 0.620 |
| age 19 strength & age 19 CGT_RT | positive | 0.05 | 0.125 | 0.06 | 0.056 |
|  | negative | -0.01 | 0.703 | **-0.07** | **0.016** |
| age 23 strength & age 23 CGT_RT | positive | 0.03 | 0.289 | 0.01 | 0.758 |
|  | negative | -0.02 | 0.564 | 0.01 | 0.744 |
| age 14 strength & age 14 SWM_BE | positive | 0.05 | 0.193 | 0.07 | 0.071 |
|  | negative | -0.02 | 0.585 | -0.03 | 0.432 |
| age 19 strength & age 19 SWM_BE | positive | 0.04 | 0.294 | 0.06 | 0.105 |
|  | negative | -0.05 | 0.180 | **-0.13** | **<0.001** |
| age 23 strength & age 23 SWM_BE | positive | **0.11** | **<0.001** | **0.09** | **0.002** |
|  | negative | **-0.09** | **0.003** | **-0.10** | **0.001** |

**Note:** RVP, Rapid Visual Information Processing; CGT, Cambridge Gambling Task; AGN, Affective Go/No-go task; RVPA , RVP Accuracy; AGN_POS, AGN mean correct latency positive; AGN_NEG, AGN mean correct latency negative; CGT_DA, CGT Delay aversion (impulsivity); CGT_RT, CGT risk-taking; SWM_BE, Strategy working memory between error.

**Table S13 Dice Coefficient (DC) of predictive networks between timepoints**

| Condition | Age | Positive |  | Negative | |
| --- | --- | --- | --- | --- | --- |
|  |  | DC | *P* value | DC | *P* value |
| Go trials | age 14 & 19 | 0.03 | 0.001 | 0.03 | <0.001 |
|  | age 14 & 23 | 0.06 | <0.001 | 0.04 | <0.001 |
|  | age 19 & 23 | 0.07 | <0.001 | 0.01 | 0.080 |
| Successful stop trials | age 14 & 19 | 0.01 | 0.235 | 0.03 | 0.001 |
|  | age 14 & 23 | 0.01 | 0.221 | 0.01 | 0.533 |
|  | age 19 & 23 | 0.01 | 0.067 | 0.00 | 1.000 |

**Table S14 Correlation between substance use and ICV at each timepoint.**

| Correlation | r value | *P* value |
| --- | --- | --- |
| age 14 ICV & age 14 alcohol | 0.03 | 0.363 |
| age 19 ICV & age 19 alcohol | 0.04 | 0.185 |
| age 23 ICV & age 23 alcohol | -0.03 | 0.342 |
| age 14 ICV & age 14 Cig+CB | 0.02 | 0.631 |
| age 19 ICV & age 19 Cig+CB | **0.13** | **<0.001** |
| age 23 ICV & age 23 Cig+CB | **0.17** | **<0.001** |
| age 19 ICV & age 19 drug | **-0.11** | **0.001** |
| age 23 ICV & age 23 drug | -0.02 | 0.556 |

Note: The bold values represent the P values that survive after FDR correction (q < 0.05).

**Table S15 Correlation between Cig+CB use and ICV across timepoints.**

| Correlation | r value | *P* value |
| --- | --- | --- |
| age 14 ICV & age 19 Cig+CB | 0.07 | 0.104 |
| age 14 ICV & age 23 Cig+CB | **0.13** | **0.007** |
| age 19 ICV & age 23 Cig+CB | **0.13** | **0.0003** |
| age 19 ICV & age 14 Cig+CB | 0.05 | 0.113 |
| age 23 ICV& age 14 Cig+CB | 0.03 | 0.352 |
| age 23 ICV & age 19 Cig+CB | **0.13** | **9.38E-05** |

Note: The bold values represent the P values that survive after FDR correction (q < 0.05).

**Table S16 Correlation between alcohol use and ICV across timepoints.**

| Correlation | r value | *P* value |
| --- | --- | --- |
| age 14 ICV & age 19 alcohol | 0.06 | 0.192 |
| age 14 ICV & age 23 alcohol | 0.03 | 0.591 |
| age 19 ICV & age 23 alcohol | 0 | 0.949 |
| age 19 ICV & age 14 alcohol | 0.02 | 0.582 |
| age 23 ICV & age 14 alcohol | -0.01 | 0.846 |
| age 23 ICV & age 19 alcohol | 0.05 | 0.12 |

Note: The bold values represent the P values that survive after FDR correction (q < 0.05).

**Table S17 Correlation between drug use and ICV across timepoints.**

| Correlation | r value | *P* value |
| --- | --- | --- |
| age 14 ICV & age 19 drug | 0.01 | 0.861 |
| age 14 ICV & age 23 drug | 0 | 0.929 |
| age 19 ICV & age 23 drug | -0.02 | 0.552 |
| age 23 ICV & age 19 drug | **-0.08** | **0.014** |

Note: The bold values represent the P values that survive after FDR correction (q < 0.05).

**Table S18 Correlation between substance use and sustained attention network at each timepoint.**

| Correlation | Predictive network | Go trials |  | Successful stop | |
| --- | --- | --- | --- | --- | --- |
|  |  | r value | *P* value | r value | *P* value |
| age 14 strength & age 14 alcohol | positive | 0.08 | 0.033 | 0.03 | 0.462 |
|  | negative | -0.07 | 0.079 | -0.03 | 0.462 |
| age 19 strength & age 19 alcohol | positive | 0.02 | 0.433 | 0.06 | 0.064 |
|  | negative | -0.03 | 0.315 | -0.03 | 0.342 |
| age 23 strength & age 23 alcohol | positive | -0.01 | 0.750 | -0.06 | 0.062 |
|  | negative | 0.05 | 0.109 | 0.07 | 0.031 |
| age 14 strength & age 14 Cig+CB | positive | -0.01 | 0.792 | 0.02 | 0.598 |
|  | negative | 0.02 | 0.552 | 0.02 | 0.646 |
| age 19 strength & age 19 Cig+CB | positive | 0.07 | 0.023 | **0.10** | **0.001** |
|  | negative | -0.07 | 0.025 | **-0.08** | **0.013** |
| age 23 strength & age 23 Cig+CB | positive | **0.12** | **<0.001** | **0.13** | **<0.001** |
|  | negative | **-0.11** | **<0.001** | **-0.11** | **0.001** |
| age 19 strength & age 19 drug | positive | -0.06 | 0.051 | **-0.12** | **<0.001** |
|  | negative | 0.06 | 0.053 | 0.05 | 0.083 |
| age 23 strength & age 23 drug | positive | -0.02 | 0.490 | -0.01 | 0.746 |
|  | negative | 0.01 | 0.795 | -0.01 | 0.752 |

Note: The bold values represent the P values that survive after FDR correction (q < 0.05).

**Table S19 Correlation between Cig+CB and sustained attention network across timepoints.**

| Correlation | Predictive network | Go trials | | Successful stop | |
| --- | --- | --- | --- | --- | --- |
|  |  | r value | *P* value | r value | *P* value |
| age 14 strength & age 19 Cig+CB | positive | 0.08 | 0.075 | 0.084 | 0.062 |
|  | negative | -0.11 | 0.019 | -0.060 | 0.187 |
| age 14 strength & age 23 Cig+CB | positive | 0.12 | 0.015 | 0.089 | 0.070 |
|  | negative | **-0.16** | **0.001** | -0.099 | 0.044 |
| age 19 strength & age 23 Cig+CB | positive | 0.06 | 0.123 | 0.085 | 0.021 |
|  | negative | -0.07 | 0.057 | -0.050 | 0.172 |
| age 19 strength & age 14 Cig+CB | positive | 0.06 | 0.078 | 0.037 | 0.240 |
|  | negative | -0.04 | 0.245 | -0.017 | 0.582 |
| age 23 strength & age 14 Cig+CB | positive | -0.01 | 0.835 | -0.022 | 0.467 |
|  | negative | 0.02 | 0.576 | 0.020 | 0.516 |
| age 23 strength & age 19 Cig+CB | positive | **0.10** | **0.002** | 0.048 | 0.138 |
|  | negative | -0.07 | 0.039 | -0.047 | 0.146 |

Note: The bold values represent the P values that survive after FDR correction (q < 0.05).

**Table S20 Correlation between alcohol use and sustained attention network across timepoints.**

| Correlation | Predictive network | Go trials |  | Successful stop | |
| --- | --- | --- | --- | --- | --- |
|  |  | r value | *P* value | r value | *P* value |
| age 14 strength & age 19 alcohol | positive | 0.01 | 0.845 | 0.03 | 0.478 |
|  | negative | -0.05 | 0.259 | -0.02 | 0.615 |
| age 14 strength & age 23 alcohol | positive | 0.06 | 0.228 | -0.02 | 0.641 |
|  | negative | 0.01 | 0.833 | -0.06 | 0.221 |
| age 19 strength & age 23 alcohol | positive | -0.08 | 0.029 | -0.02 | 0.648 |
|  | negative | 0.06 | 0.111 | 0.00 | 0.997 |
| age 19 strength & age 14 alcohol | positive | 0.05 | 0.150 | 0.03 | 0.421 |
|  | negative | 0.00 | 0.948 | 0.02 | 0.448 |
| age 23 strength & age 14 alcohol | positive | 0.01 | 0.653 | 0.00 | 0.960 |
|  | negative | 0.00 | 0.904 | 0.03 | 0.369 |
| age 23 strength & age 19 alcohol | positive | 0.04 | 0.176 | 0.06 | 0.075 |
|  | negative | -0.07 | 0.025 | -0.02 | 0.538 |

Note: The bold values represent the P values that survive after FDR correction (q < 0.05).

**Table S21 Correlation between drug use and sustained attention network across timepoints.**

| Correlation | Predictive network | Go trials | | Successful stop | |
| --- | --- | --- | --- | --- | --- |
|  |  | r value | *P* value | r value | *P* value |
| age 14 strength & age 19 drugs | positive | 0.00 | 0.918 | -0.036 | 0.429 |
|  | negative | 0.09 | 0.049 | -0.025 | 0.580 |
| age 14 strength & age 23 drugs | positive | -0.05 | 0.297 | -0.028 | 0.570 |
|  | negative | 0.07 | 0.179 | 0.076 | 0.121 |
| age 19 strength & age 23 drugs | positive | -0.02 | 0.618 | -0.013 | 0.723 |
|  | negative | 0.03 | 0.431 | 0.023 | 0.525 |
| age 23 strength & age 19 drugs | positive | -0.07 | 0.039 | **-0.091** | **0.005** |
|  | negative | 0.07 | 0.036 | **0.087** | **0.007** |

Note: The bold values represent the P values that survive after FDR correction (q < 0.05).

**Table S22 Cig+CB use between different sustained attention groups (at age 14) at each timepoint.**

|  | Low SA | High SA | t value | *P* value |
| --- | --- | --- | --- | --- |
| ICV |  |  |  |  |
| Age 14 | -0.151 | -0.138 | 0.739 | 0.462 |
| Age 19 | 0.0826 | -0.269 | -2.38 | **0.020** |
| Age 23 | 0.218 | -0.322 | -3.32 | **0.001** |
| Positive Network strength | | | | |
| Age 14 | -0.139 | -0.138 | 0.0556 | 0.956 |
| Age 19 | 0.166 | -0.294 | -2.78 | **0.007** |
| Age 23 | 0.109 | -0.351 | -3.06 | **0.003** |
| Negative network strength | | | | |
| Age 14 | -0.129 | -0.138 | -0.691 | 0.491 |
| Age 19 | 0.206 | -0.24 | -2.35 | **0.021** |
| Age 23 | 0.182 | -0.269 | -2.76 | **0.007** |

**Table S23 Cig+CB use between different working memory groups (at age 14) at each timepoint.**

|  | Low WM | High WM | t value | *P* value |
| --- | --- | --- | --- | --- |
| Age 14 | -0.0713 | -0.0771 | 0.111 | 0.912 |
| Age 19 | 0.0534 | -0.121 | 1.65 | 0.0995 |
| Age 23 | 0.0188 | -0.131 | 1.43 | 0.154 |

**Table S24 Prediction for intra-individual coefficient of variation (ICV) across timepoints controlling for ICV at age 14.**

| Predictive models | Predictive networks | Go trials |  | Successful stop trials | |
| --- | --- | --- | --- | --- | --- |
|  |  | r value | *P* value | r value | *P* value |
| Age 14 predicts Age 19 | positive | **0.10** | **0.028** | **0.08** | **0.036** |
|  | negative | 0.06 | 0.119 | **0.10** | **0.012** |
|  | combined | **0.08** | **0.047** | **0.11** | **0.009** |
| Age 14 predicts Age 23 | positive | **0.11** | **0.013** | **0.11** | **0.005** |
|  | negative | 0.04 | 0.187 | **0.13** | **0.005** |
|  | combined | 0.08 | 0.056 | **0.13** | **0.017** |
| Age 19 predicts Age 23 | positive | **0.22** | **<0.001** | **0.18** | **<0.001** |
|  | negative | **0.19** | **<0.001** | **0.18** | **<0.001** |
|  | combined | **0.22** | **<0.001** | **0.17** | **<0.001** |

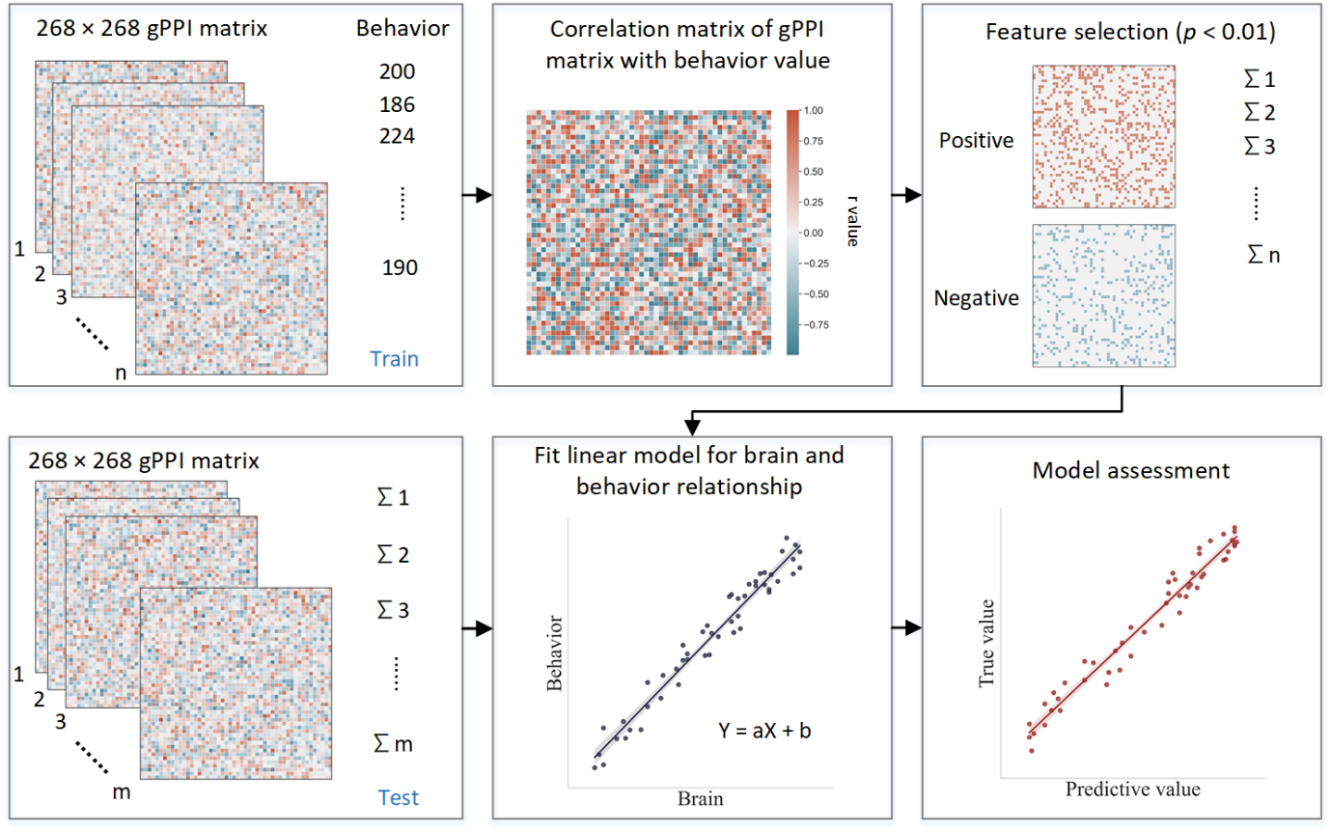
 **Fig. S1 Schematic of connectome-based predictive modeling.** i) Feature selection. The correlation between each edge in the gPPI matrix and the behavioral phenotype is calculated while controlling for several covariates in the training set. These covariates include age, gender, mean framewise displacement (mean FD), scan sites, and mode-centered PDS (only for age 14). The r value with the associated p-value for each edge is obtained using partial correlation, and a threshold of *P =*0.01 is used to select the edges. Positively or negatively correlated edges are regarded as positive or negative networks. Network strength is then calculated by summing the selected edges in the gPPI matrix for both positive and negative networks, as well as by subtracting the strength of the negative network from the strength of the positive network to obtain the combined network strength. ii) Model building. Linear models are constructed between the network strength of the positive, negative, combined network, and behavioral phenotype in the training set. The network strength is then calculated for each subject in the testing set and input into the predictive model along with covariates to yield a predicted behavioral phenotype (e.g., predicted ICV) for each network. iii) Model validation. The predictive performance is evaluated by calculating the correlation between predicted and observed values.

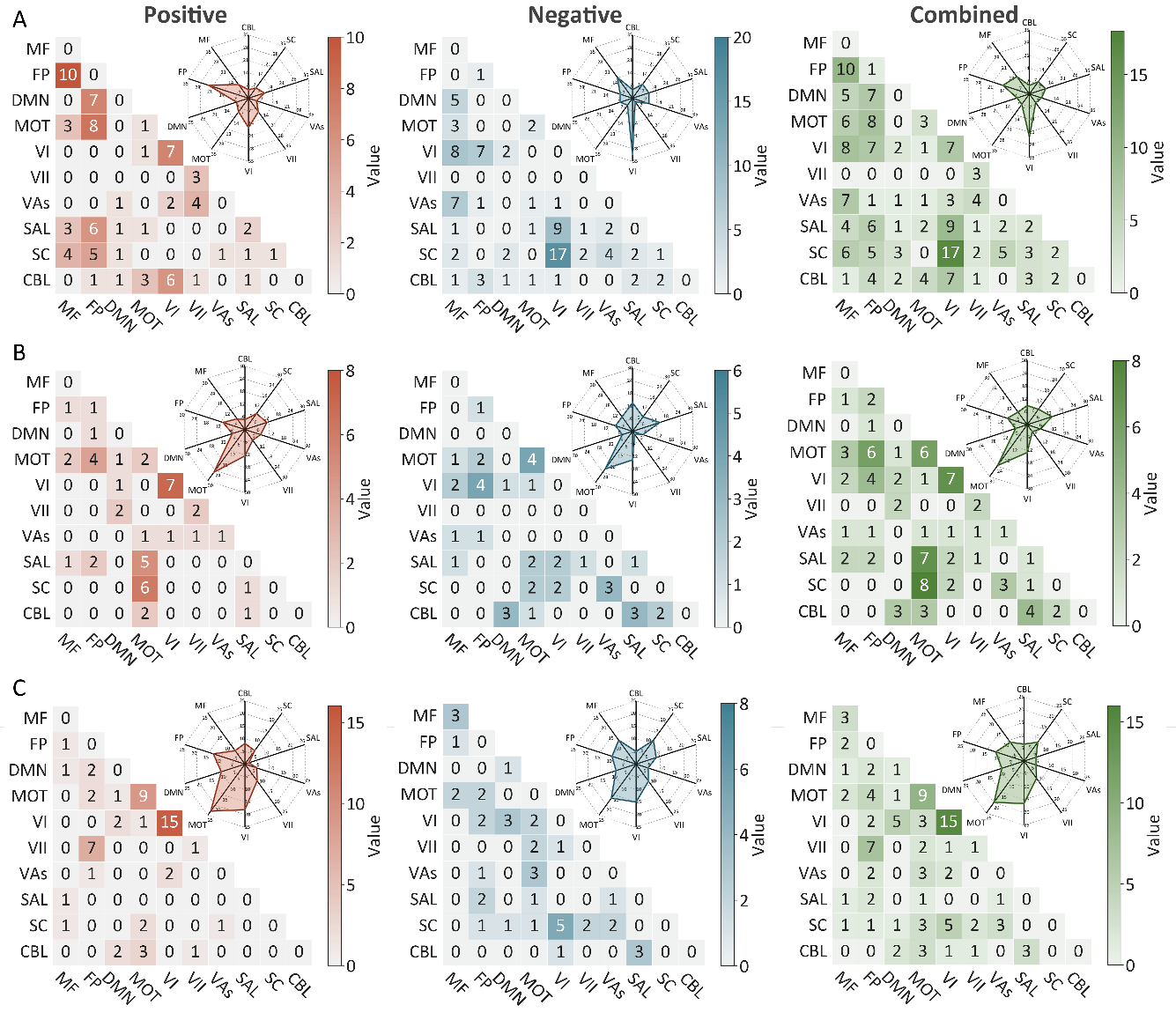

**Fig. S2 The predictive networks predicting intra-individual coefficient of variation (ICV)** **per timepoint derived from Go trials.** Heatmap of networks for ICV, predictive networks are those with non-zero values at **(A)** age 14 **(B)** age 19 **(C)** age 23. Each heatmap cell shows the number of edges between or within functional networks. Radar plots show the proportion of the functional networks involved in predictive networks. The edges depicted above are those selected in at least 95% of cross validation folds. Red, blue, and green represent positive, negative, and combined networks separately. MF, medial frontal; FP, frontoparietal; DMN, default mode; MOT, motor; VI, Visual I Network; VII, Visual II Network; VAs, Visual Association Network; SAL, salience; SC, Subcortical; CBL, Cerebellar. R/L, right/left hemisphere.

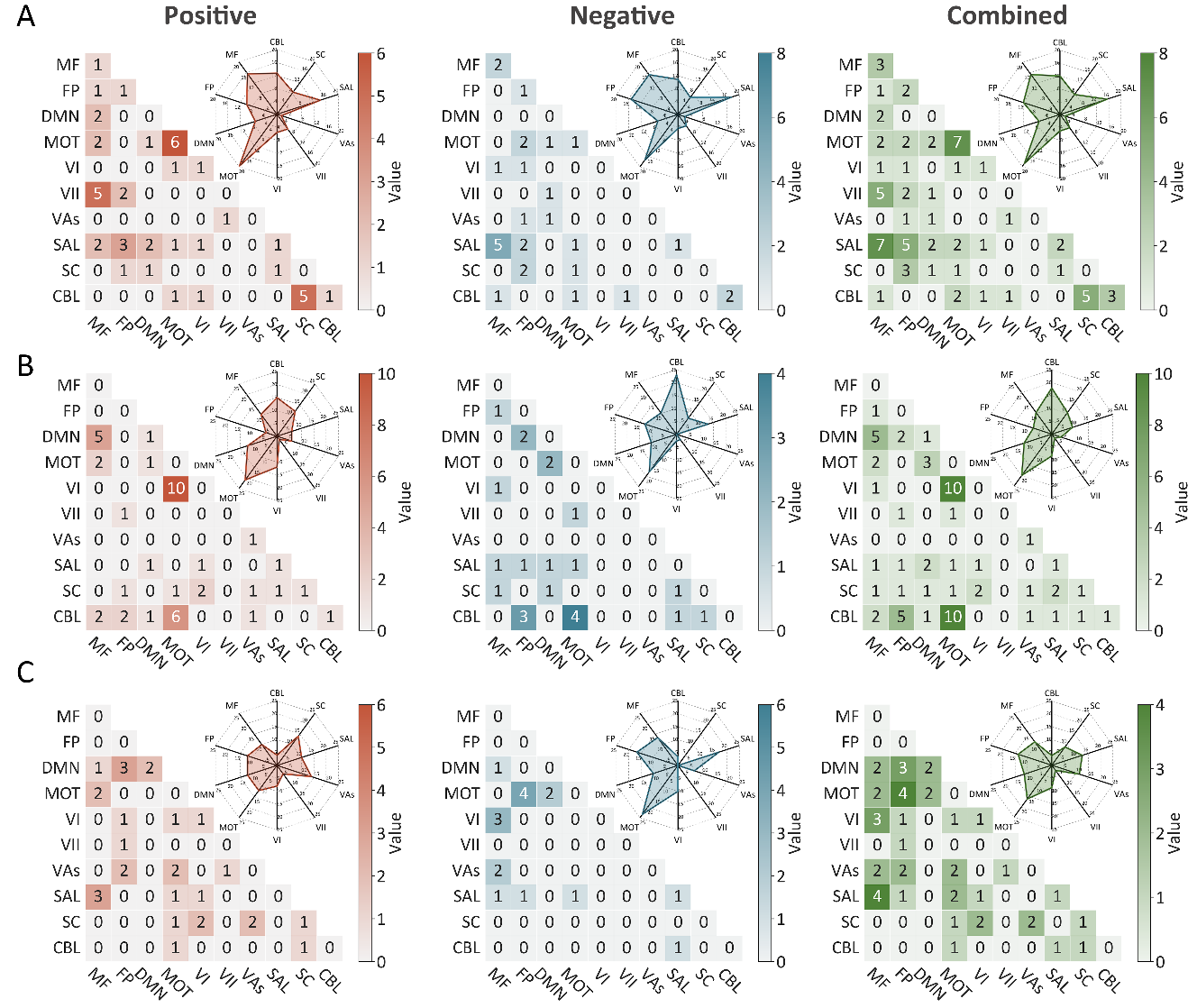

**Fig. S3 The predictive networks predicting intra-individual coefficient of variation (ICV)** **per timepoint derived from Successful stop trials.** Heatmap of networks for ICV, predictive networks are those with non-zero values at **(A)** age 14 **(B)** age 19 and **(C)** age 23. Each heatmap cell shows the number of edges between or within functional networks. Radar plots show the proportion of the functional networks involved in predictive networks. The edges depicted above are those selected in at least 95% of cross validation folds. Red, blue, and green represent positive, negative, and combined networks separately. MF, medial frontal; FP, frontoparietal; DMN, default mode; MOT, motor; VI, Visual I Network; VII, Visual II Network; VAs, Visual Association Network; SAL, salience; SC, Subcortical; CBL, Cerebellar. R/L, right/left hemisphere.

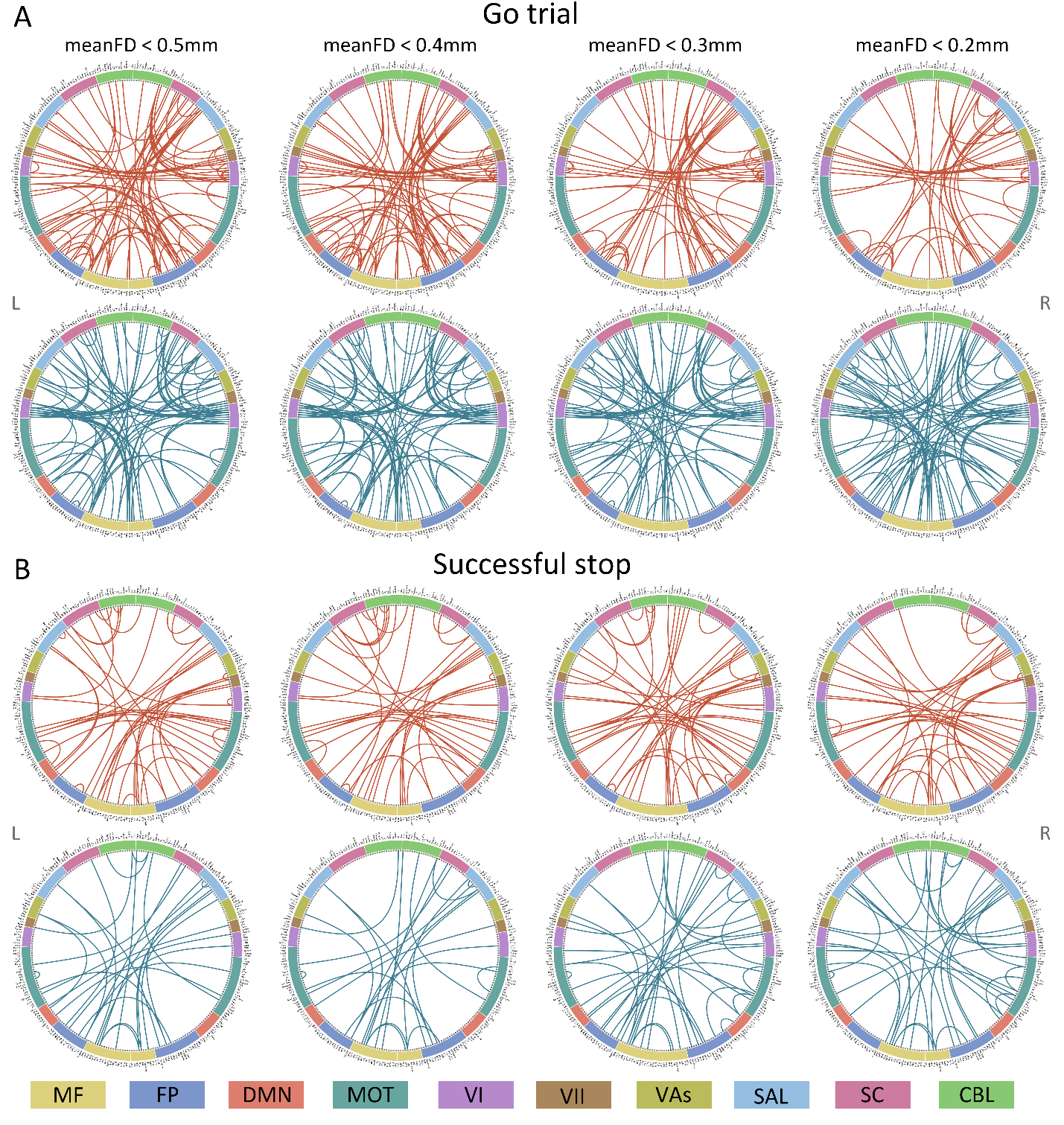

**Fig. S4 Connectome of positive and negative networks predicting ICV at age 14 with mean framewise displacement (meanFD) from 0.2mm to 0.5mm.** Red and blue lines represent positive, negative, and combined networks separately. MF, Medial frontal; FP, Frontoparietal; DMN, Default mode; MOT, Motor; VI, Visual I; VII, Visual II; VAs, Visual association; SAL, Salience; SC, Subcortical; CBL, Cerebellar. R/L, right/left hemisphere.

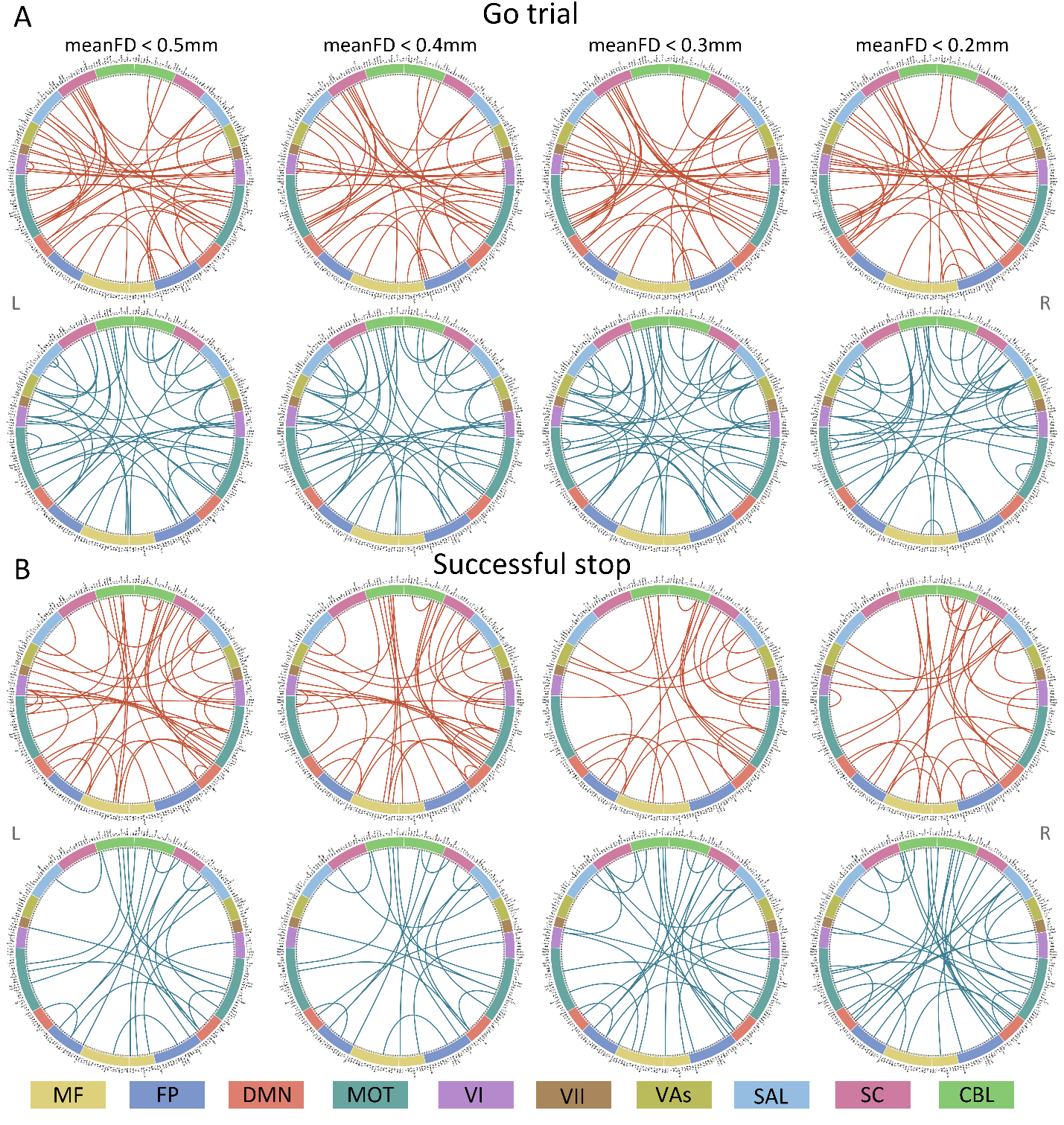

**Fig. S5 Connectome of positive and negative networks predicting ICV at age 19 with mean framewise displacement (meanFD) from 0.2 mm to 0.5mm.** Red and blue lines represent positive, negative, and combined networks separately. MF, Medial frontal; FP, Frontoparietal; DMN, Default mode; MOT, Motor; VI, Visual I; VII, Visual II; VAs, Visual association; SAL, Salience; SC, Subcortical; CBL, Cerebellar. R/L, right/left hemisphere.

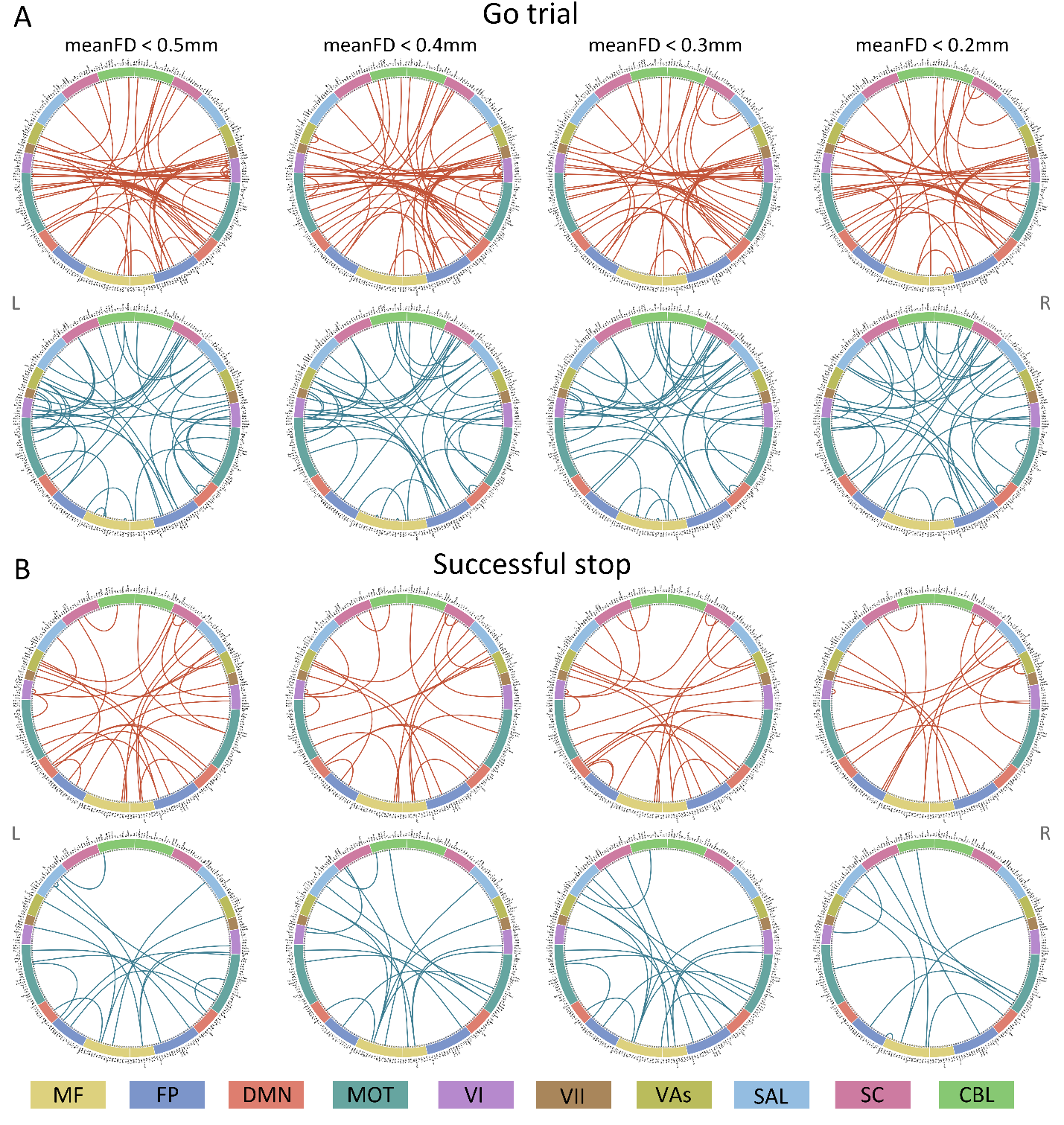
 **Fig. S6 Connectome of positive and negative networks predicting ICV at age 23 with mean framewise displacement (meanFD) from 0.2mm to 0.5mm.** Red and blue lines represent positive, negative, and combined networks separately. MF, Medial frontal; FP, Frontoparietal; DMN, Default mode; MOT, Motor; VI, Visual I; VII, Visual II; VAs, Visual association; SAL, Salience; SC, Subcortical; CBL, Cerebellar. R/L, right/left hemisphere.

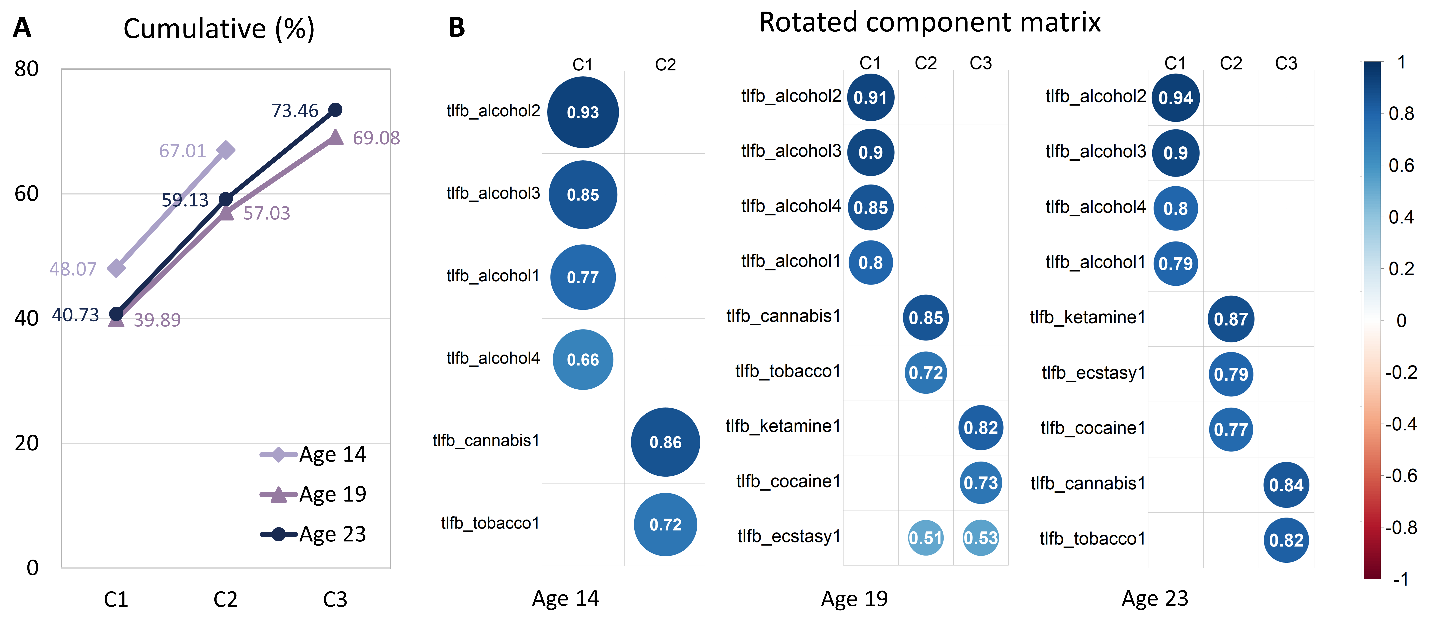

**Fig. S7 Exploratory factor analysis of TLFB at each timepoint.** (A) Total variance explained for exploratory factor analysis of TLFB items. (B) Rotated component matrix for exploratory factor analysis. Extraction Method: Principal Component Analysis. Rotation Method: Varimax with Kaiser Normalization.

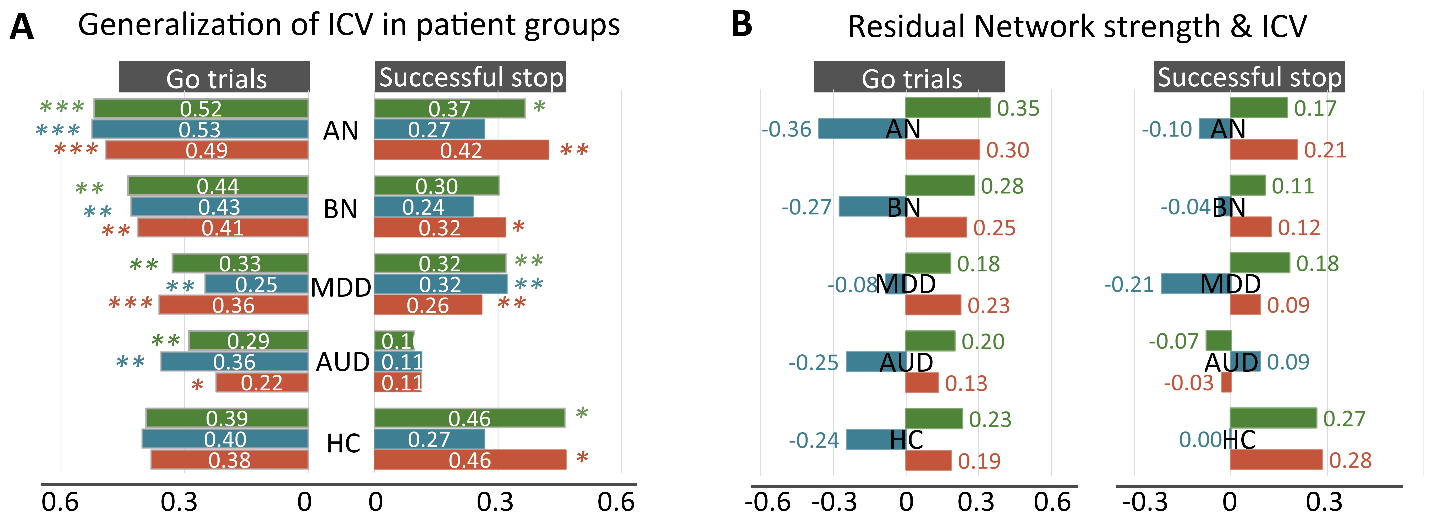

**Fig. S8 Generalization in subgroups in STRATIFY.** (A) Predictive performance in distinct patient groups in STRATIFY derived from Go and Successful stop trials. (B) The correlation between network strength and ICV across patient cohorts in STRATIFY derived from Go and Successful stop trials. AUD, alcohol use disorder; MDD, major depression disorder; BN, bulimia nervosa; AN, anorexia nervosa; HC, healthy controls. *, *P <* 0.05; **, *P <* 0.01; ***, *P* < 0.001.

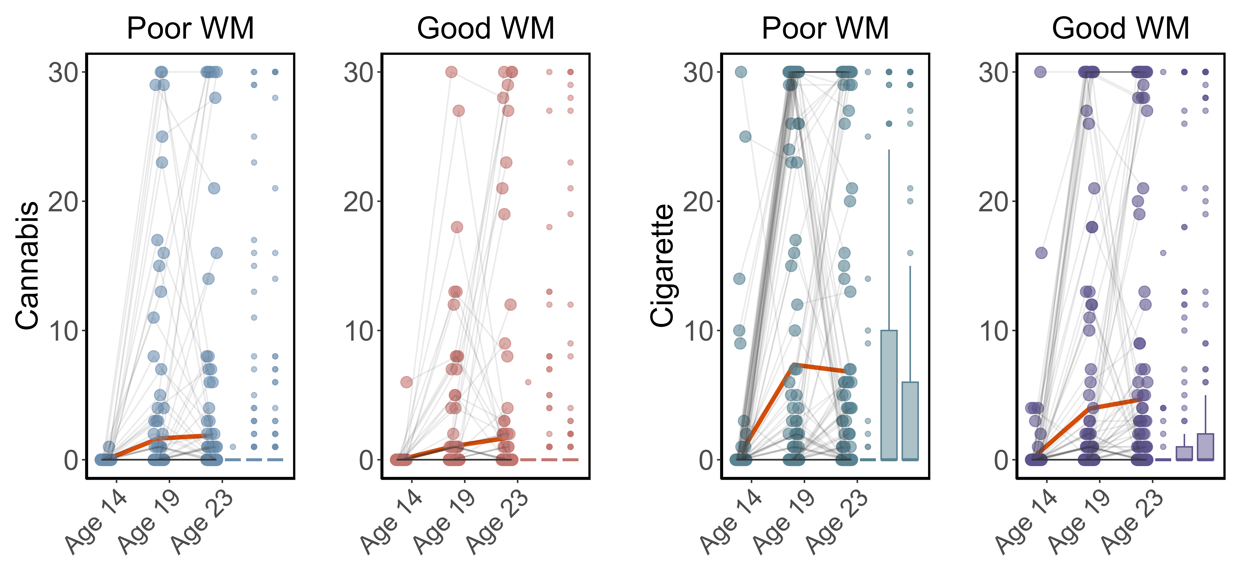

**Fig. S9 Cigarette and cannabis score in Timeline Followback change in individuals with good working memory (Good WM) and poor working memory (Poor WM) from ages 14 to 23.** Participants were categorized into five equal groups based on the performance of Strategy working memory task at age 14.

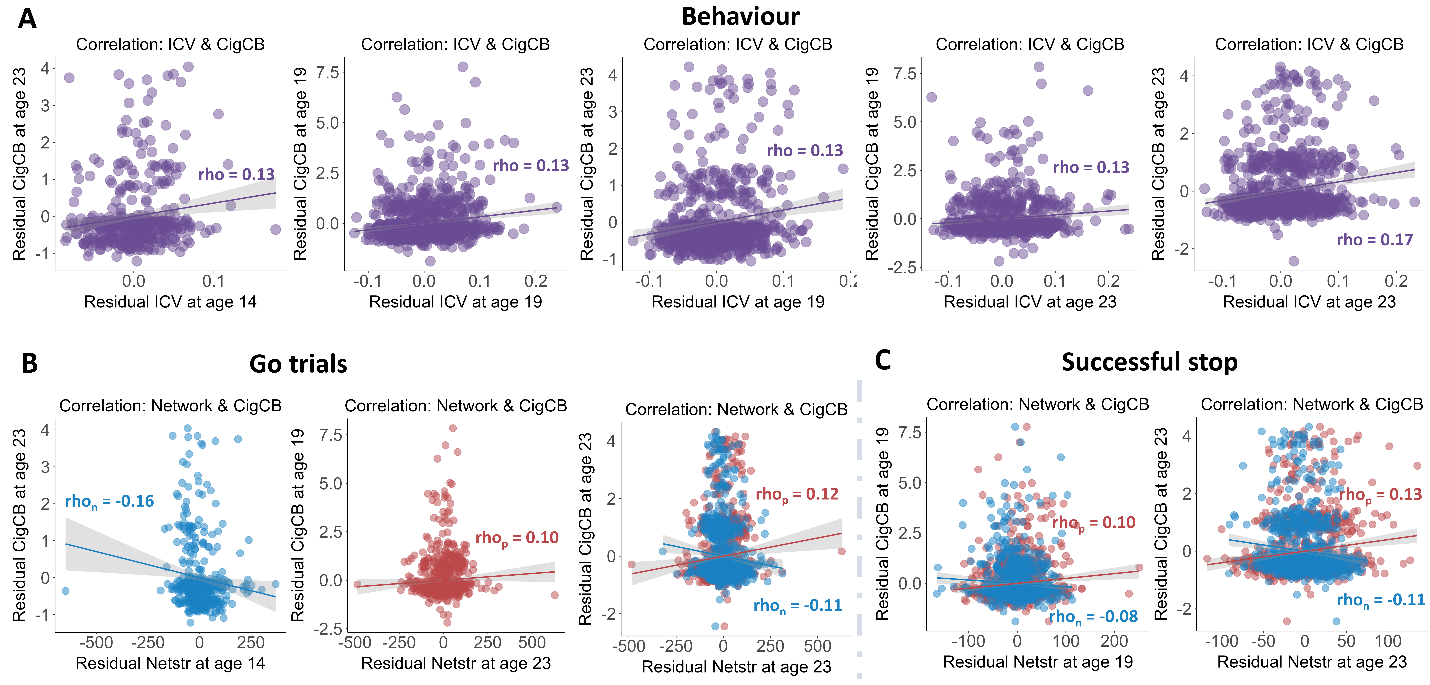

**Fig. S10 Significant correlations between sustained attention and substance use across timepoints (FDR correction, q<0.05).** (A) Correlations between the intra-individual coefficient of variation (ICV) and Cigarette and cannabis use (Cig+CB) across timepoints. Correlations between sustained attention network strength and Cig+CB across timepoints (B) derived from Go trials and (C) derived from Successful stop trials. Rho_p_: r value between network strength of the positive network. Rho_n_: r value between network strength of the negative network.
